# APOE knockout attenuates vascular graft fibrosis by limiting profibrotic macrophage formation through low-density lipoprotein receptor related protein 1

**DOI:** 10.1101/2025.07.22.666232

**Authors:** Jiayin Fu, Meng Zhao, Jing Zhao, Shaofei Wu, Jiahui Wu, Xulin Hong, He Huang, Guosheng Fu, Shengjie Xu

## Abstract

Vascular graft fibrosis can cause a decrease in cellular infiltration and capillary ingrowth in vascular walls and vascular stiffening. As such, there are still no vascular grafts that can be used in blood vessels where their diameters are less than 6 mm in patients. Although various approaches have been evaluated to mitigate implant-associated fibrosis, effective treatments remain quite limited. In this study, we demonstrated that APOE was significantly increased during vascular regeneration after graft implantation *in vivo*. APOE knockout (KO) increased compliance of regenerated aortas and reduced extracellular matrix (ECM) deposition in adventitia of the regenerated aortas. Using single cell RNA sequencing (scRNA-seq), a subset of profibrotic macrophages was found to be involved in graft fibrosis and APOE KO limited the formation of profibrotic macrophage formation during vascular regeneration. The interaction between APOE and low-density lipoprotein receptor related protein 1 (LRP1) partially mediated fibrotic differentiation of the macrophages. Profibrotic macrophages promoted graft fibrosis mainly through secretion of insulin-like growth factor-1 (IGF-1) that could support proliferation of fibroblasts. Finally, we showed that APOE knockdown *in vivo* using adeno-associated virus (AAV) improved the compliance of regenerated aortas and reduced ECM deposited in the adventitial areas by limiting formation of profibrotic macrophages. Collectively, these data indicate that APOE promotes the profibrotic transition of macrophages partially through LRP1, and the profibrotic macrophages increase the proliferation of fibroblasts via IGF-1. Inhibition of APOE by AAV can alleviate graft fibrosis occurring during vascular regeneration.

## Introduction

Fibrosis is characterized by an excessive accumulation of connective tissues that mainly consists of fibroblasts and extracellular matrix (ECM) proteins, such as collagens and fibronectin, following long-term and unresolved chronic inflammation^1-3^. Because of chronic inflammations caused by biomaterials or devices after their implantation *in vivo* (foreign body reactions), fibrosis can occur surrounding implants. Over time, fibrous tissues formed surrounding implants can act as barriers to hinder physical, electrical, chemical and nutrient communications^4-6^, ultimately resulting in implant dysfunction. With a dramatic increase in prosthesis implanted in recent years, the implant-associated fibrosis gradually becomes a major concern of clinicians and patients.

Small-diameter vascular grafts (internal diameters <6 mm) fabricated with polycaprolactone (PCL) using electrospinning are first reported by Walpoth *et al.* in year 2008. The electrospun PCL grafts show fast endothelial coverage and homogeneous neointima formation after implantation *in vivo*^7^. However, cellular infiltration and capillary ingrowth dramatically decrease in vascular walls 18 month after implantation *in vivo*^8^. This is probably because thick fibrous capsules surrounding the grafts block mass transfer to vascular walls. In addition, graft fibrosis can also lead to vascular stiffening, which increases blood pressure^9^ and causes dysfunction of endothelium due to increased shear stress of blood flow to vascular walls^10^. As a result, even though it has been 17 years since their first development, electrospun PCL vascular grafts still cannot be successfully used in blood vessels where their diameters are less than 6 mm.

To mitigate implant-associated fibrosis caused by foreign body reactions, various approaches have been evaluated, including coating biomaterials with anti-inflammation drugs^11^, hydrogel coatings^12^ or adhesives^13^, and manipulation of physical properties of implanted materials, such as stiffness^14^, size^15^, and surface roughness^16^. However, effective treatments for implant fibrosis remain quite limited. The main reason is probably that most of these studies focus on implants themself but ignore host reactions in response to implants. For small-diameter vascular grafts, although researchers have focused on acceleration of endothelium recovery and inhibition of intimal hyperplasia to prevent failure of vascular grafts^17-22^, there are currently quite few studies to mitigate graft fibrosis after implantation, which, however, is equally important for their successful application in patients.

Apolipoprotein E (APOE), a glycoprotein of 35 kDa, plays a key role in lipid metabolism by clearance of APOE-rich lipoprotein particles via low density lipoprotein receptor (LDLR) family mainly in liver^23,24^. APOE also mediates the uptake of lipid nanoparticles by hepatocytes that highly express LDLR^25^ and cells in bone marrows^26^. In addition, APOE is implicated in unresolvable inflammation, such as Alzheimer’s disease, atherosclerosis^27^, and osteoarthritis^28^. The most recent data suggests that APOE may have a close relationship with fibrosis. One recent study shows that alveolar macrophages express APOE in bleomycin-induced lung fibrosis, which mediates phagocytosis of collagens by alveolar macrophages to resolve fibrosis^29^. Another recent study highlights the role of APOE in promoting survival of monocyte-derived alveolar macrophages able to improve lung fibrosis^30^. Both studies identify APOE as a key factor during cure of lung fibrosis.

Fibroblasts are the main effector cells for deposition and remodeling of ECM, and their functions are regulated by macrophages during fibrosis^31^. Fibroblasts respond to macrophage-derived signals to proliferate and deposit ECM^32,33^. With advancement of single cell RNA sequencing (scRNA-seq), a high heterogeneity of macrophages during chronic inflammation is revealed and a subset of profibrotic macrophages is identified to have function of promoting fibrosis^34^. These profibrotic macrophages reside in fibrotic niches and activate fibroblasts via secretion of various cytokines^35^, such as platelet derived growth factor (PDGF)-AA^36^ and resistin-like molecule α^37^. For vascular regeneration after graft implantation *in vivo*, monocytes are recruited into the grafts where they differentiate into macrophages^38^. If macrophages with profibrotic phenotypes are involved in graft fibrosis and how their profibrotic differentiation is regulated are largely unknown.

Here we demonstrated that APOE expression significantly increased during vascular regeneration after graft implantation *in vivo*. APOE knockout (KO) increased compliance of regenerated aortas and reduced ECM deposition in adventitia of the regenerated aortas. Using scRNA-seq, a subset of profibrotic macrophages were found to be involved in graft fibrosis, and APOE KO limited the formation of profibrotic macrophage formation during vascular regeneration. The interaction between APOE and low-density lipoprotein receptor related protein 1 (LRP1) partially mediated the profibrotic differentiation of macrophages, which promoted graft fibrosis through insulin-like growth factor-1 (IGF-1) that enhanced fibroblast proliferation. Finally, we showed that APOE knockdown *in vivo* using adeno-associated virus (AAV) greatly attenuated graft fibrosis occurring during vascular regeneration.

## Results and discussion

### Increased APOE expression during vascular regeneration after graft implantation *in vivo*

To understand how different types of cells were orchestrated during vascular regeneration after graft implantation *in vivo*, scRNA-seq was performed. Seven different types of cells were identified in the regenerated and native aortas according to marker genes of each cell type (**Supplementary Fig. 1a and b**). A dynamic change in percentages of different types of cells over time was observed (**Supplementary Fig. 1c and d**). Crosstalk between different cell types was analyzed further. The results showed that there was much more crosstalk between different cell types in the regenerated aortas than in the native aortas (**Supplementary Fig. 2a**). In detail, in the native aortas, crosstalk between cells was realized mainly via amyloid-beta precursor protein (APP) signaling and interactions between collagens and integrins. In contrast, crosstalk between cells relied on prosaposin (PSAP) signaling, apolipoprotein (APOE) signaling, APP signaling, and interactions between fibronectin and integrins in the regenerated aortas (**Supplementary Fig. 2b**). These results indicate that there is rewiring of pathways by which crosstalk between cells are realized during vascular regeneration after graft implantation *in vivo*.

Since APOE signaling was among the top three pathways that mediated crosstalk between cells in the regenerated aortas (**Supplementary Fig. 2b**), we then examined the expression of APOE in native and regenerated aortas at different timepoints during vascular regeneration after graft implantation *in vivo*. The scRNA-seq data showed an increase in APOE expression over time (**Fig. 1a**). The endothelial cells (ECs), macrophages and neurons had a higher expression of APOE than any other cell type (**Fig. 1b**). There was a gradual increase in APOE expression for both ECs and neurons during the period of implantation. For macrophages, the expression of APOE increased from Day 14 to Day 30 and decreased from Day 30 to Day 90. The other types of cells, such as B cells, fibroblasts, smooth muscle cells (SMCs) and T cells, were not active in APOE expression (**Fig. 1c**). Immunofluorescence staining showed that there were few APOE positive cells in the native aortas, but the regenerated aortas were intensively stained with APOE (**Fig. 1d**). Both western blot (WB) and enzyme linked immunosorbent assay (ELISA) showed similar results with low and high APOE expression in native aortas and regenerated aortas, respectively (**Fig. 1e and f**). All these results indicate that the expression of APOE dramatically increased after vascular graft implantation *in vivo*. APOE expression is also increased in osteoarthritis, which promotes proteoglycan degeneration of cartilages^28^. Moreover, APOE mediates uptake of lipid nanoparticles by livers^26^ and bone marrows^25,39^. It seems that APOE participates in chronic inflammation and nanoparticle uptake *in vivo*. However, the role that APOE plays during vascular regeneration after graft implantation is unclear.

**Fig. 1.**
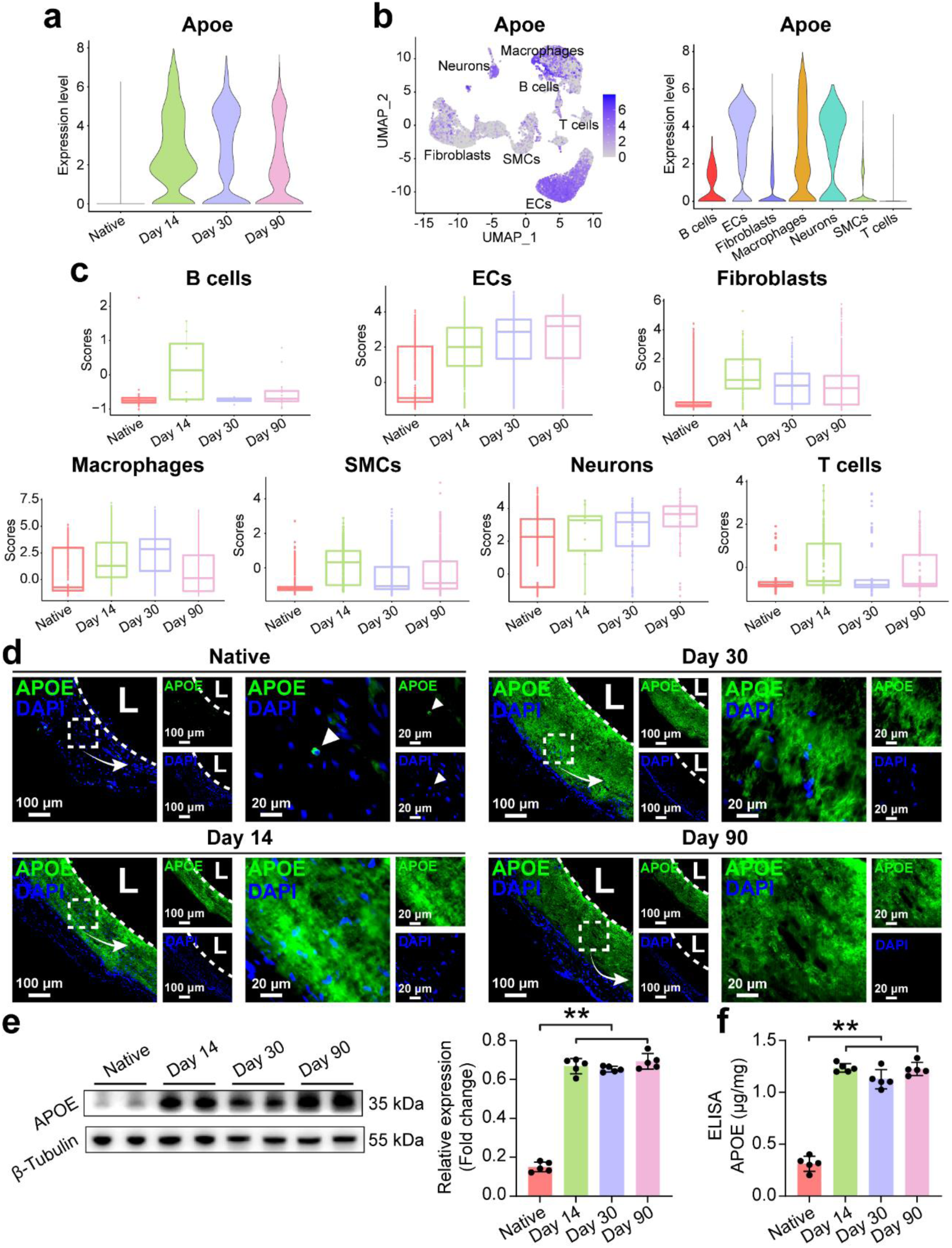
Increased APOE expression during vascular regeneration after graft implantation *in vivo*. (a) Violin plots of Apoe expression in native and regenerated aortas across different timepoints post graft implantation. (b) UMAP and violin plots of Apoe expression in different cell clusters in native and regenerated aortas. (c) Box plots of Apoe expression in different cell clusters in native and regenerated aortas across different timepoints post graft implantation. (d) Immunofluorescence staining of APOE in native and regenerated aortas across different timepoints post graft implantation. L indicates lumens. Arrow heads indicate cells positively stained by APOE. (e) WB analysis of APOE levels in native and regenerated aortas across different timepoints post graft implantation and quantification of APOE levels. ** indicates p < 0.01, Tukey’s post-hoc test. For each timepoint, five different samples from five different animals were used (n=5). (f) Quantification of APOE by ELISA. ** indicates p < 0.01, Tukey’s post-hoc test. For each timepoint, five different samples from five different animals were used (n=5).

### Increased compliance of regenerated aortas in APOE knockout (KO) rats

To investigate the role of APOE during vascular regeneration after graft implantation, APOE knockout (KO, Apoe^-/-^) rats were used in this study. The performances of vascular grafts implanted in wide type (WT) and APOE KO rats were evaluated by ultrasound imaging. As shown in **Fig. 2a and b**, grafts implanted in both WT and APOE KO rats could stay patent on Day 30 and Day 90, respectively. Regenerated aortas in APOE KO rats showed higher resistance index (RI) and pulsatility index (PI) than those in WT rats (**Fig. 2c and d**). Furthermore, the regenerated aortas in WT rats had no vascular wall movements, while the regenerated aortas in APOE KO rats showed vascular wall movements like those observed in the native aortas (**Fig. 2e**). This result suggests that the regenerated aortas in APOE KO rats can dilate and contract with blood pressure cycles. The regenerated aortas in APOE KO rats had a compliance of 6.15% and 9.17% on Day 30 and 90 respectively, whereas the regenerated aortas in WT rats only showed a compliance of 2.00% on Day 30 and 1.61% on Day 90 (**Fig. 2f**). These data demonstrate that APOE KO increases the compliance of the regenerated aortas.

**Fig. 2.**
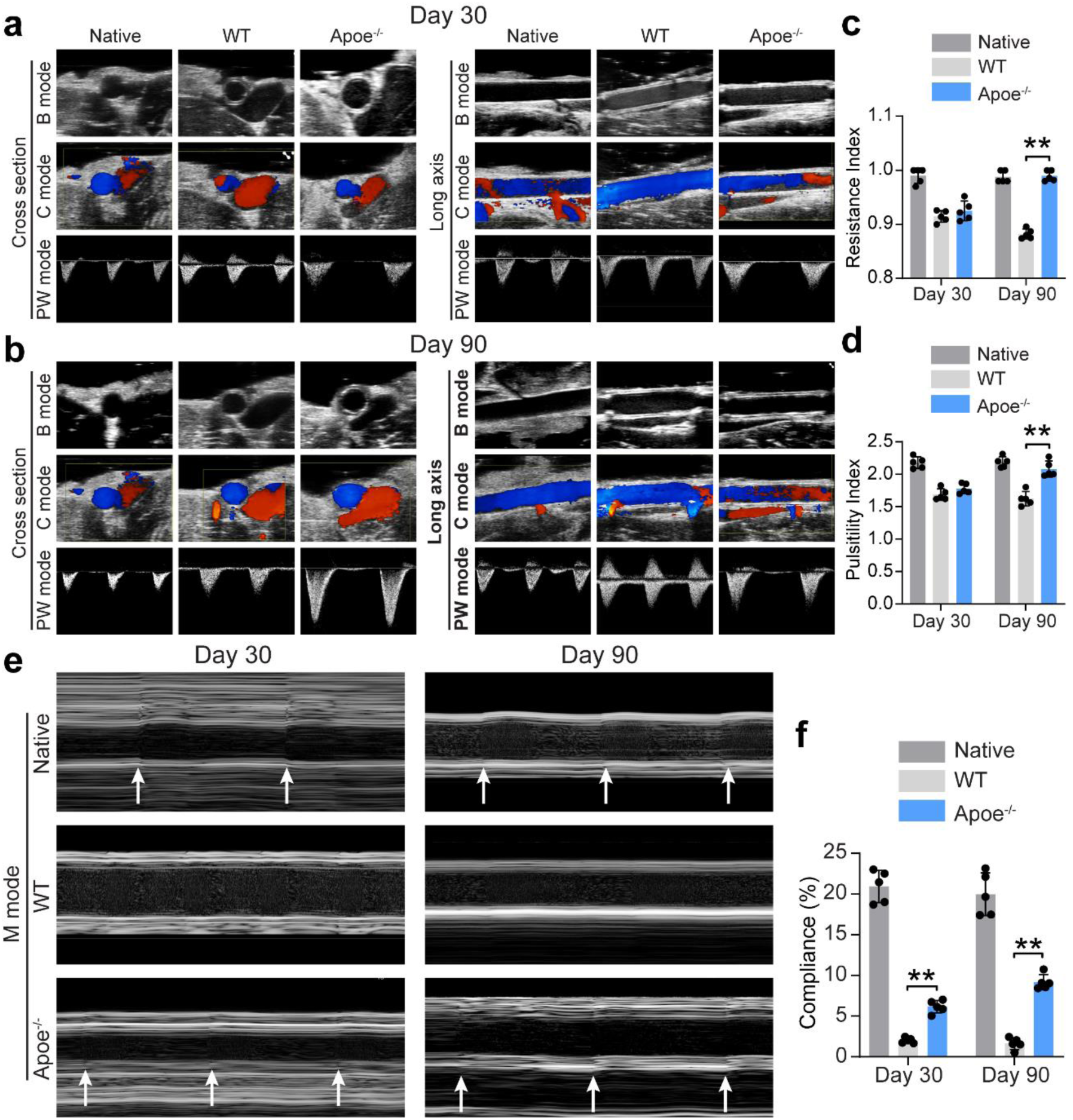
APOE KO increasing compliance of regenerated aortas. Ultrasound imaging of native and regenerated aortas in WT and Apoe^-/-^ rats at Day 30 (a) and Day 90 (b). Quantification of RI (c) and PI (d) of native and regenerated aortas in WT and Apoe^-/-^ rats at Day 30 and Day 90. ** indicates p < 0.01, Tukey’s post-hoc test. For each timepoint and each group, five different images from five different animals were used (n=5). (e) M mode images of ultrasound of native and regenerated aortas in WT and Apoe^-/-^ rats at Day 30 and Day 90. Arrow heads indicate movement of vascular walls. (f) Quantification of compliance of native and regenerated aortas in WT and Apoe^-/-^ rats at Day 30 and Day 90. ** indicates p < 0.01, Tukey’s post-hoc test. For each timepoint and each group, five different images from five different animals were used (n=5).

### Reduction in ECM deposition in adventitia of regenerated aortas after APOE KO

Next, the histology of regenerated aortas in WT and APOE KO rats was evaluated. As shown in **Fig. 3a**, the regenerated aortas in WT rats were reddish, while the regenerated aortas in APOE KO rats were pale. Immunoflurescene staining confirmed KO of gene Apoe, as no positive staining of APOE could be observed in the regenerated aortas in APOE KO rats (**Fig. 3b**). H&E staining showed thicker adventitia in the regenerated aortas in WT rats, as compared to those in APOE KO rats. Masson’s trichrome (MTC) showed that APOE KO resulted in less collagen deposition in adventitial areas of regenerated aortas (**Fig. 3c and d**). ECM proteins, such as collagen I (COL I), collagen III (COL III) and lumican (LUM), were expressed less in adventitial areas of regenerated aortas in WT rats than in APOE KO rats (**Fig. 3e-f**).

**Fig. 3.**
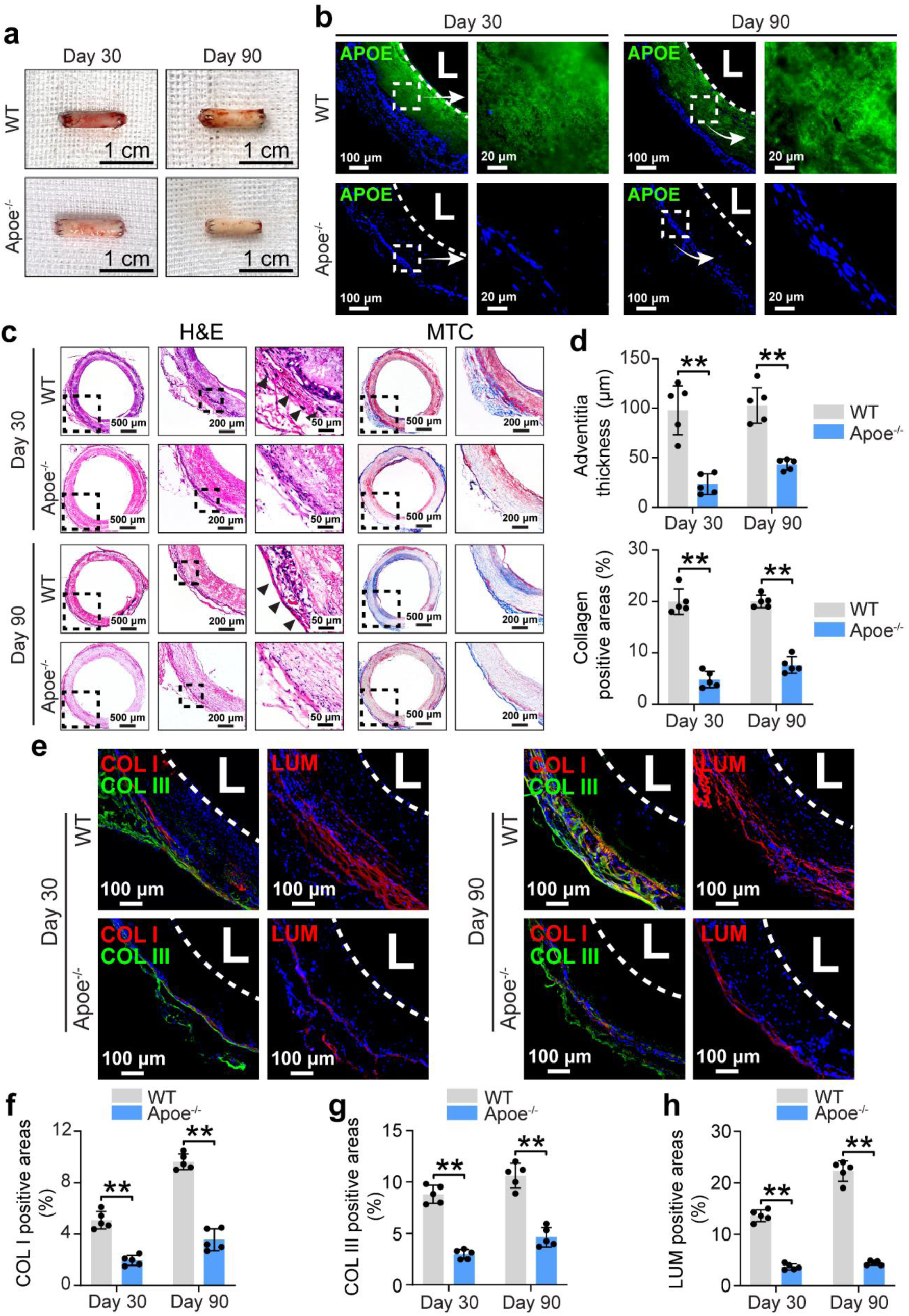
APOE KO reducing ECM deposition in adventitial areas of regenerated aortas. (a) Representative images of regenerated aortas harvested from WT and Apoe^-/-^ rats at Day 30 and Day 90. (b) Immunofluorescence staining of APOE in regenerated aortas from WT and Apoe^-/-^ rats at Day 30 and Day 90. L indicates lumens. (c) H&E and MTC staining of regenerated aortas from WT and Apoe^-/-^ rats at Day 30 and Day 90. (d) Quantification of adventitia thickness and collagen positive areas in regenerated aortas from WT and Apoe^-/-^ rats at Day 30 and Day 90. ** indicates p < 0.01, Tukey’s post-hoc test. For each timepoint and each group, five different samples from five different animals were used (n=5). (e) Immunofluorescence staining of COL I, COL III and LUM in regenerated aortas from WT and Apoe^-/-^ rats at Day 30 and Day 90. L indicates lumens. Quantification of COL I positive areas (f), COL III positive areas (g), and LUM positive areas (h) in regenerated aortas from WT and Apoe^-/-^ rats at Day 30 and Day 90. ** indicates p < 0.01, Tukey’s post-hoc test. For each timepoint and each group, five different samples from five different animals were used (n=5).

ScRNA-seq data confirmed these results. When combining the scRNA-seq data of regenerated aortas from WT and APOE KO rats 30 days post implantation, seven different types of cells could be identified (**Supplementary Fig. 3a-c**). Among them, the fibroblasts had the highest ECM scores (**Supplementary Fig. 3d)**, indicating that fibroblasts were the main effector of fibrosis. Compared with the fibroblasts from APOE KO rats, the fibroblasts from WT rats were more highly enriched in ECM-related genes (**Supplementary Fig. 3e**) and had higher ECM scores (**Supplementary Fig. 3f and g**). The scRNA-seq data of regenerated aortas from WT and APOE KO rats 90 days post implantation showed similar results, with fibroblasts from WT rats expressing higher ECM-related genes than those from APOE KO rats (**Supplementary Fig. 4**).

Finally, a subcutaneous implantation model was used as well to verify that APOE KO could reduce ECM deposition. As shown in **Supplementary Fig. 5 a and c,** thicker fibrous capsules were formed after electrospun PCL sheets were subcutaneously implanted. In contrast, the fibrous capsules formed in APOE KO rats were thin. Meanwhile, MTC staining showed less collagen deposition surrounding PCL sheets in APOE KO rats than in WT rats (**Supplementary Fig. 5 a and d**). Immunofluorescence staining also showed that there were more COL I, CoL III, and LUM deposition surrounding PCL sheets in WT rats than in APOE KO rats on Day 30 and Day 90, respectively (**Supplementary Fig. 5b and e-g**). Taken together, APOE KO can reduce fibrosis of regenerated aortas as well as implanted biomaterials.

### Profibrotic macrophages involved in vascular regeneration

Then, we investigated how APOE affected fibrosis of regenerated aortas. Here, we first supposed that APOE could stimulate ECM production by fibroblasts directly. However, there was no increase in the expressions of COL I and FN by fibroblasts after APOE stimulation (**Supplementary Fig. 6**). Since macrophages are master regulators of fibrosis^31,40-42^, we then analyzed the scRNA-seq data of macrophages to see if APOE could affect phenotypes of macrophages. Macrophages involved in vascular regeneration could be further divided into 9 subgroups (**Fig. 4a**), indicating a high heterogeneity of macrophages. Among them, the cluster 2 (C2) macrophages highly expressed Ctsd, Spp1, Gpnmb, Lgals3, and Fabp5 (**Fig. 4b and d**), which accounted for 20% to 30% of the total macrophages at different timepoints of vascular regeneration. However, there were few such macrophages in native aortas (**Fig. 4c**). Immunofluorescence staining confirmed that there were macrophages co-expressed CD68 and Cathepsin D (CTSD) at different timepoints of vascular regeneration (**Fig. 4e**). The double positive cells also existed in the native aortas, but they were quite few (**Supplementary Fig. 7**). The protein levels of CTSD and secreted phosphoprotein 1 (SPP1) increased from Day 14 to Day 90 post graft implantation but decreased on Day 90 (**Fig. 4f**). The *in vitro* study also showed that the expression of CTSD and SPP1 increased with the increase of APOE when macrophages were seeded on PCL scaffolds (**Supplementary Fig. 8a and b**). Ctsd, Spp1, Gpnmb, Lgals3, and Fabp5 are signature genes of profibrotic macrophages^31,43,44^. Therefore, we demonstrate here that profibrotic macrophages are involved in vascular regeneration after graft implantation *in vivo*. Since APOE significantly increased during vascular regeneration (**Fig. 1**), it was highly possible that APOE in vascular regeneration niches induced profibrotic phenotypes of macrophages.

**Fig. 4.**
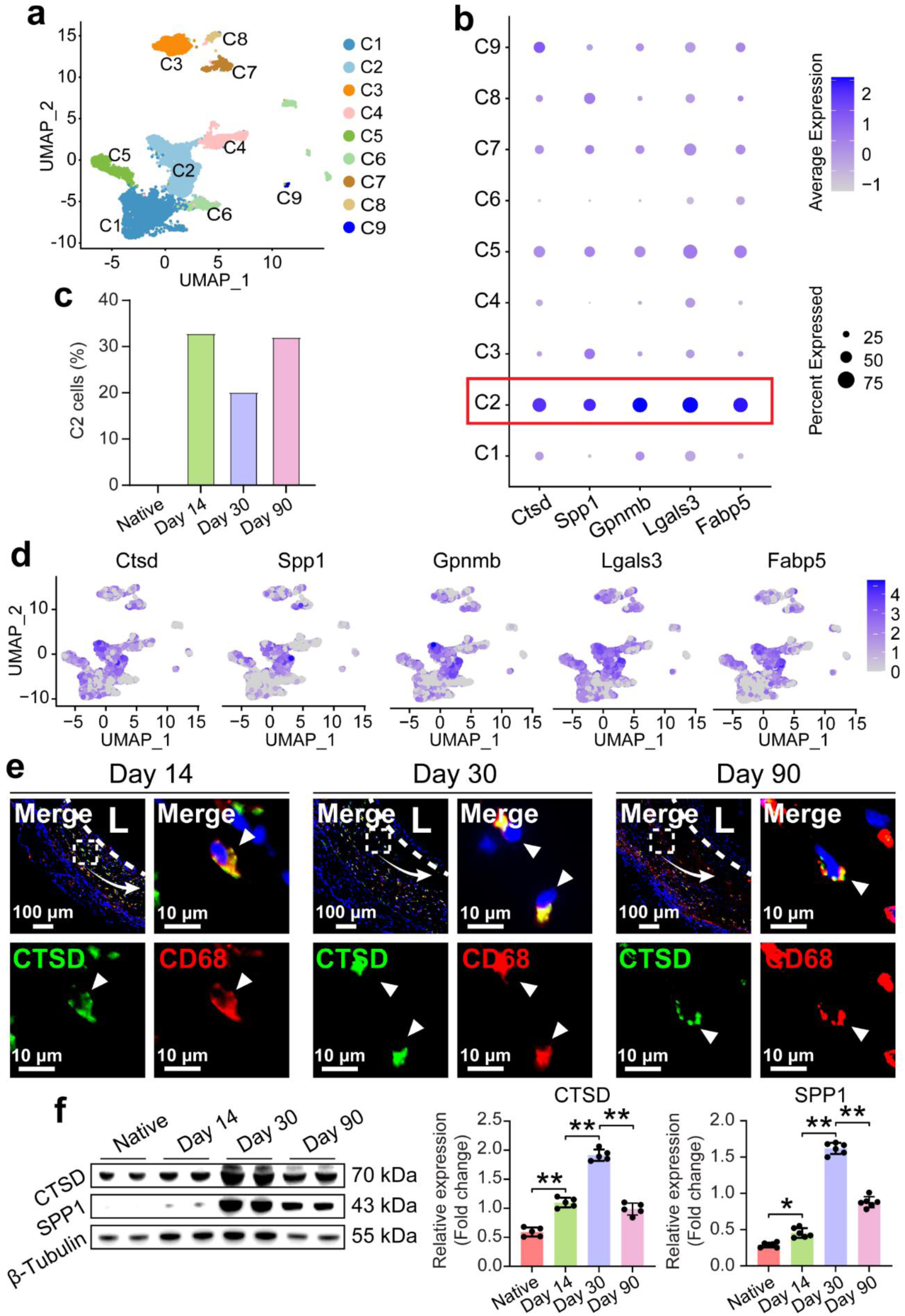
Profibrotic macrophages being involved in vascular regeneration. (a) UMAP of macrophages in native aortas and regenerated aortas. (b) Dot plots of marker genes for subgroup of macrophages. (c) Percentage of cluster 2 (C2) macrophages in native aortas and regenerated aortas across different timepoints post graft implantation. (d) Expression of Ctsd, Spp1, Gpnmb, Lgals3, and Fabp5 in UMAP of macrophages in native aortas and regenerated aortas. (e) Immunofluorescence staining of CD68 and CTSD in regenerated aortas across different timepoints post graft implantation. L indicates lumens. Arrow heads indicate positively stained cells. (f) WB analysis of levels of CTSD and SPP1 in native and regenerated aortas across different timepoints post graft implantation and quantification of their levels. ** indicates p < 0.01, Tukey’s post-hoc test. For each timepoint and each group, five different samples from five different animals were quantified (n=5).

### APOE KO limiting profibrotic macrophage formation during vascular regeneration

Next, we evaluated the effect of APOE on differentiation of macrophages towards a profibrotic state. Interestingly, scores of profibrotic macrophage genes (Ctsd, Spp1, Gpnmb, Lgals3, and Fabp5) greatly reduced in Apoe^-/-^ rats on Day 30 and the percentage of C2 macrophages dropped from 32.38% to 6.59% (**Fig. 5a**). Although the gene set scores increased in Apoe^-/-^ rats on Day 90, the percentage of C2 macrophages (19.33%) in Apoe^-/-^ rats was still lower than that in WT rats (32.07%, **Fig. 5b**). Immunofluorescence staining revealed more CD68^+^CTSD^+^ cells in WT rats than those in Apoe^-/-^ rats on both Day 30 and Day 90 (**Fig. 5c and d**). WB results also showed that the protein levels of CTSD and SPP1 decreased following APOE KO (**Fig. 5e and f**). *In vitro* data showed that the expressions of CTSD and SPP1 were inhibited when culturing macrophages from Apoe^-/-^ rats on PCL scaffolds (**Fig. 5g and h**). Meanwhile, positive staining of APOE, CTSD and SPP1 could be observed in WT macrophages but not in APOE KO macrophages. These results highly support the assumption that APOE can promote profibrotic phenotype transition of macrophages. One previous study shows that APOE can promote survival of macrophages by increasing macrophage colony-stimulating factor secretion in lung fibrosis^30^. In fact, a higher percentage of macrophages could be observed in regenerated aortas from APOE KO rats than from WT rats on Day 30 and Day 90 (**Supplementary Fig. 3c and Supplementary Fig. 4c**). Therefore, we believe APOE is necessary for macrophage profibrotic transition rather than for their survival during vascular regeneration after graft implantation *in vivo*.

**Fig. 5.**
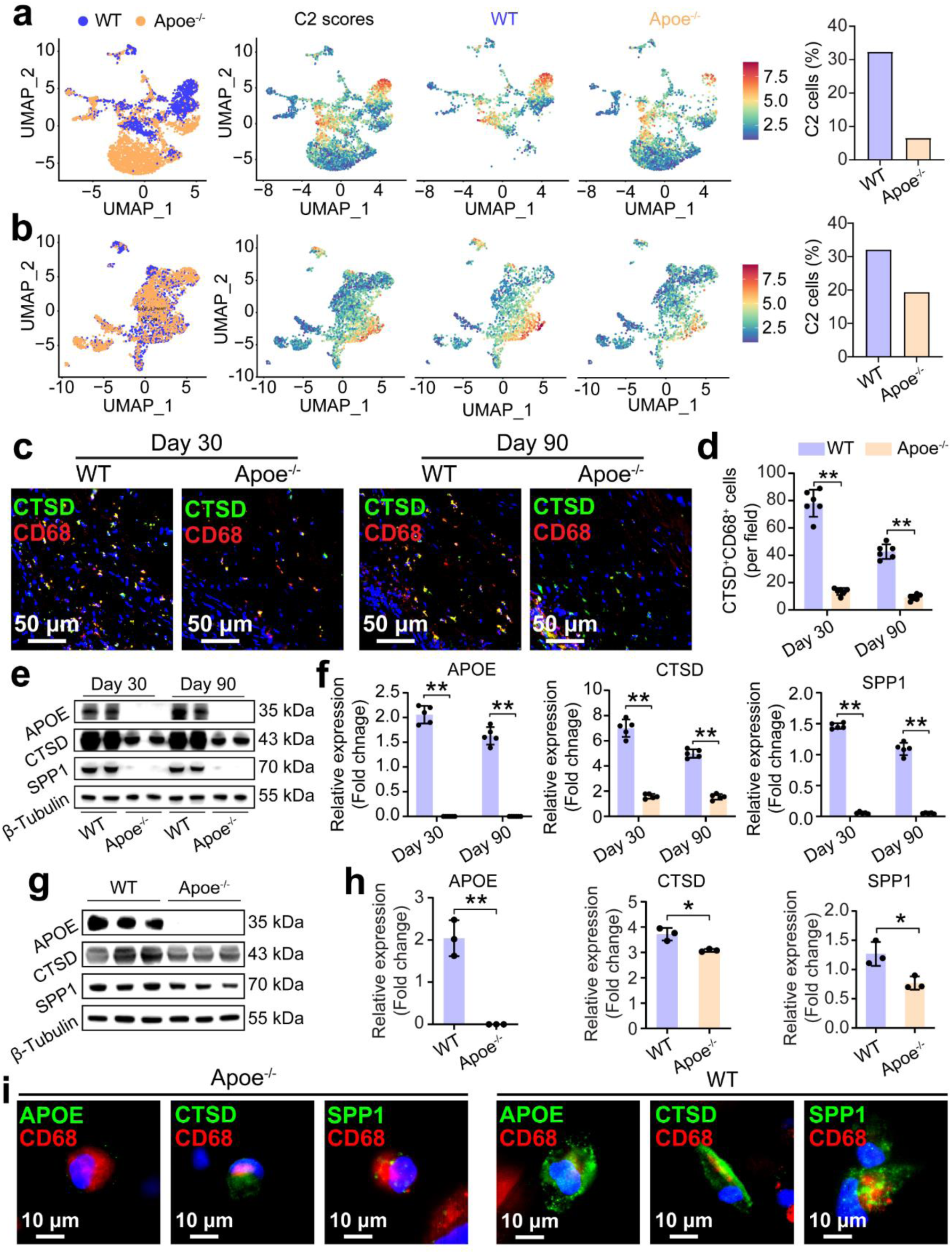
APOE KO reducing profibrotic macrophages during vascular regeneration. UMAP of macrophages in regenerated aortas post graft implantation in WT and Apoe^-/-^ rats, heatmap of C2 scores in the UMAP of macrophages, and the percentage of C2 cells in the macrophages at Day 30 (a) and Day 90 (b). (c) Immunofluorescence staining of CD68 and CTSD in regenerated aortas post graft implantation in WT and Apoe^-/-^ rats at Day 30 and Day 90. (d) Quantification of CD68 and CTSD double positive cells. ** indicates p < 0.01, Tukey’s post-hoc test. For each timepoint and each group, five different samples from five different animals were quantified (n=5). (e) WB analysis of levels of APOE, CTSD and SPP1 in regenerated aortas post graft implantation in WT and Apoe^-/-^ rats at Day 30 and Day 90. (f) Quantification of levels of APOE, CTSD and SPP1. ** indicates p < 0.01, Tukey’s post-hoc test. For each timepoint and each group, five different samples from five different animals were quantified (n=5). (g) WB analysis of levels of APOE, CTSD and SPP1 in macrophages from WT and Apoe^-/-^ rats after their culture on PCL scaffolds for 48 hours. (h) Quantification of levels of APOE, CTSD and SPP1 in the macrophages. ** indicates p < 0.01, unpaired t test. For each timepoint and each group, three different samples were quantified (n=3). (i) Immunofluorescence staining of APOE and CD68, CTSD and CD68, SPP1 and CD68 in macrophages from WT and Apoe^-/-^ rats after their culture on PCL scaffolds for 48 hours.

### Identification of LRP1 as a receptor of APOE for macrophage profibrotic differentiation

Subsequently, immunoprecipitation (IP) following mass spectrometry (MS) was performed to identify potential receptors of APOE for macrophage profibrotic differentiation. Seven different types of receptors that could possibly interact with APOE were identified by MS (**Fig. 6a**). Co-immunoprecipitation (Co-IP) confirmed that there was direct interaction between APOE and low-density lipoprotein receptor-related protein 1 (LRP1) (**Fig. 6b**). Co-localization of LRP1 with CD68 in regenerated aortas at different timepoints, indicating that macrophages involved in vascular regeneration expressed this receptor (**Fig. 6c**). Confocal imaging showed co-localization of LRP1 with APOE in macrophages cultured on PCL scaffolds *in vitro* (**Fig. 6d**), which further confirms the interaction between LRP1 and APOE. Adenovirus (ADV) encoding shRNA targeting Lrp1 (ADV-shRNA(Lrp1)) was used to knock down the expression of LRP1 in macrophages. As shown in **Fig. 6e and f**, ADV-shRNA(Lrp1) treatment successfully downregulated the expression of LRP1 as compared to the macrophages treated with ADV encoding scramble shRNA (ADV-shRNA(NC)). Although ADV treatment didn’t affect the expression of APOE (**Fig. 6e and g**), the protein levels of CTSD and SPP1 in macrophages cultured on PCL scaffolds were inhibited by ADV-shRNA(Lrp1) (**Fig. 6e, h and i**). Flowcytometry showed that ADV-shRNA(Lrp1) treatment decreased CTSD^+^ macrophages from 57% to 44% (**Fig. 6j and k**). These results reveal that LRP1 partially mediates the effect of APOE on promoting macrophage profibrotic differentiation. LRP1 belongs to low-density lipoprotein receptor (LDLR) family and plays an important role in preservation of cardiovascular homeostasis^45^. Deletion of macrophage LRP1 increases atherogenesis due to decreased efferocytosis by macrophages^46^. LRP1 also plays an important role in nervous system disease and lung fibrosis. In Alzheimer’s disease, LRP1 mediates endocytosis of tau protein and the downregulation of LRP1 can reduce tau uptake by neurons^47^. In bleomycin-induced lung fibrosis, LRP1 is required for collagen phagocytosis by macrophages to resolve lung fibrosis^29^. These studies suggest an important role of LRP1 in mediation of endocytosis. It is highly possible that the APOE/LRP1 pathway mediates the endocytosis of degradation product of PCL grafts used in this study, which then promotes the profibrotic transition of macrophages.

**Fig. 6.**
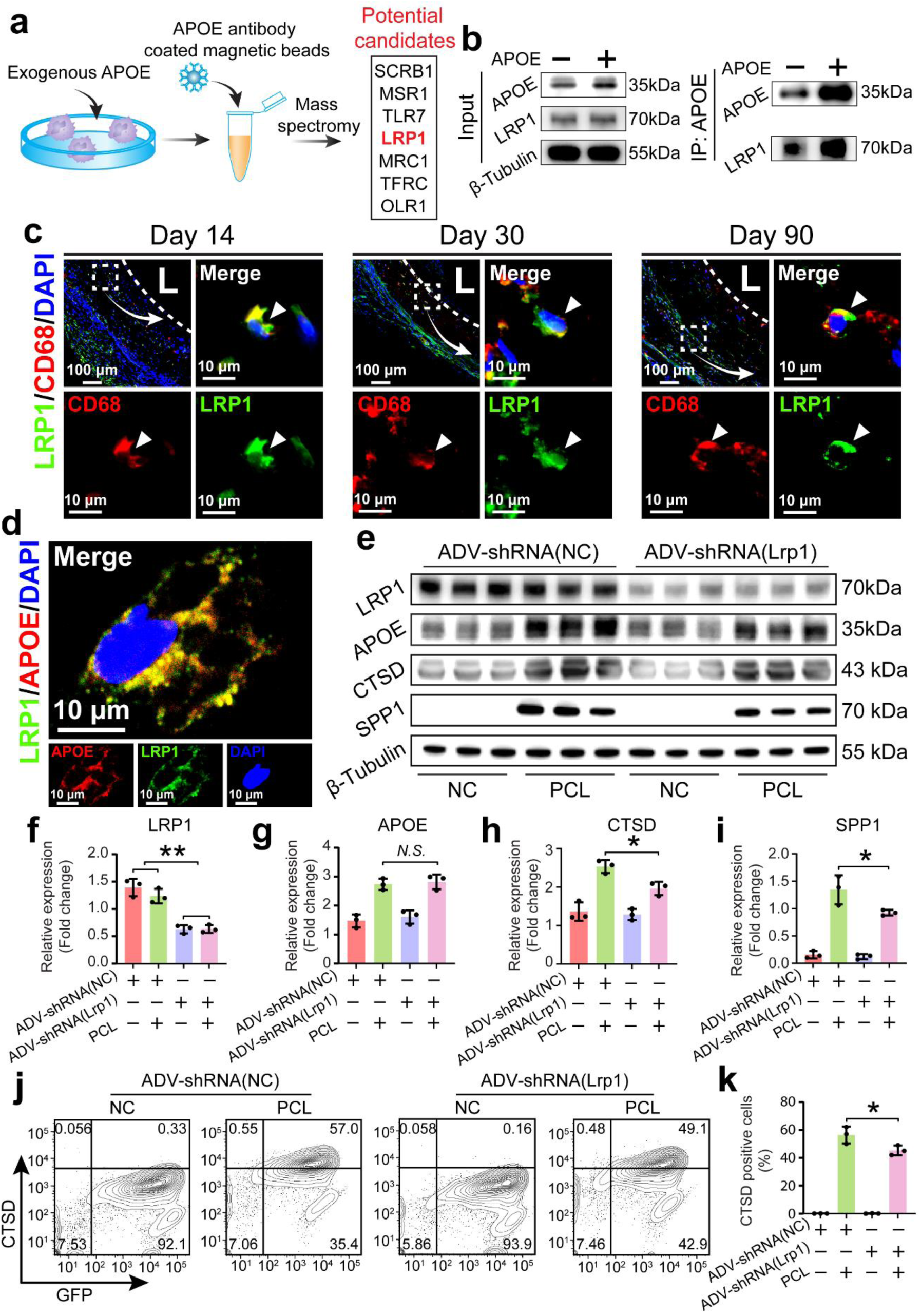
APOE/LRP1 interaction promoting profibrotic transition of macrophages during vascular regeneration. (a) Immunoprecipitation (IP) following mass spectrometry (MS) to identify the potential receptors of APOE on surfaces of macrophages. (b) Co-immunoprecipitation (Co-IP) to confirm interaction between APOE and LRP1. (c) Immunofluorescence staining of CD68 and LRP1 in regenerated aortas post graft implantation across different timepoints. (d) Immunofluorescence staining of APOE and LRP1 in macrophages from WT rats after 48 hours of culture on PCL scaffolds. (e) WB analysis of levels of LRP1, APOE, CTSD and SPP1 in macrophages cultured on tissue culture plates (negative control, NC) or PCL scaffolds (PCL) for 48 hours prior to treatment with shRNA ADV-shRNA(NC) or ADV-shRNA(Lrp1) for 24 hours. (h) Quantification of levels of LRP1, APOE, CTSD and SPP1 in macrophages. * indicates p < 0.05, ** indicates p < 0.01, Tukey’s post-hoc test. For each timepoint and each group, three different samples were quantified (n=3). (e) Flow cytometry of CTSD positive cells in macrophages cultured on tissue culture plates (negative control, NC) or PCL scaffolds (PCL) for 48 hours prior to treatment with shRNA ADV-shRNA(NC) or ADV-shRNA(Lrp1) for 24 hours and quantification of percentage of CTSD positive cells. * indicates p < 0.05, Tukey’s post-hoc test. For each timepoint and each group, three different samples were quantified (n=3).

### IGF-1 secreted by profibrotic macrophages increasing fibroblast proliferation

Finally, we explored how profibrotic macrophages promoted graft fibrosis during vascular regeneration. Igf1, encoding insulin-like growth factor-1 (IGF-1), was highly expressed in C2 macrophages as compared to the other subgroups of macrophages on both Day 30 and Day 90 (**Fig. 7a**). Since a lower percentage of the C2 macrophages in APOE KO rats (**Fig. 5**) than in WT rats, the Igf1 expression score in Apoe^-/-^ rats was lower than that in WT rats (**Fig. 7b**), as expected. The concentration of IGF-1 in regenerated aortas in Apoe^-/-^ rats was also lower than that in WT rats, as determined by the enzyme linked immunosorbent assay (ELISA, **Fig. 7c**). Cell cycle genes (Ccnd1, Ccnd2, and Ccnd3) were then used to evaluate proliferation status of fibroblasts. As shown in **Fig. 7d**, the cell cycle gene scores of fibroblasts in WT rats were significantly higher than those in Apoe^-/-^ rats on Day 30. A similar result also could be observed in regenerated aortas on Day 90 (**Fig. 7e**). In consistence with the scRNA-seq data, Ki67 positive cells in the regenerated aortas from WT rats were higher than those from Apoe^-/-^ rats (**Fig. 7f and g**). *In vitro* cell culture experiments showed that there was a lower concentration of IGF-1 in cell culture media when Apoe^-/-^ macrophages were seeded on PCL scaffolds as compared to WT macrophages (**Fig. 7h**). Medium conditioned by WT macrophages but not Apoe^-/-^ macrophages increased proliferation of fibroblasts. IGF-1 blocking antibody (Ab) treatment inhibited the proliferation of fibroblasts stimulated by WT macrophage conditioned medium but had no effect on proliferation of fibroblasts treated with the medium conditioned by Apoe^-/-^ macrophages (**Fig. 7i**). All these results reveal that IGF-1 secreted by profibrotic macrophages increases the proliferation of fibroblasts, thereby promoting fibrosis during vascular regeneration after graft implantation *in vivo*. The level of IGF-1 is elevated in pulmonary fibrosis^48,49^ and IGF-1 is required for fibroblast activation to promote lung fibrosis^36,50^. Here, we demonstrate that macrophage-derived IGF-1 also plays an important role in vascular graft fibrosis.

**Fig. 7.**
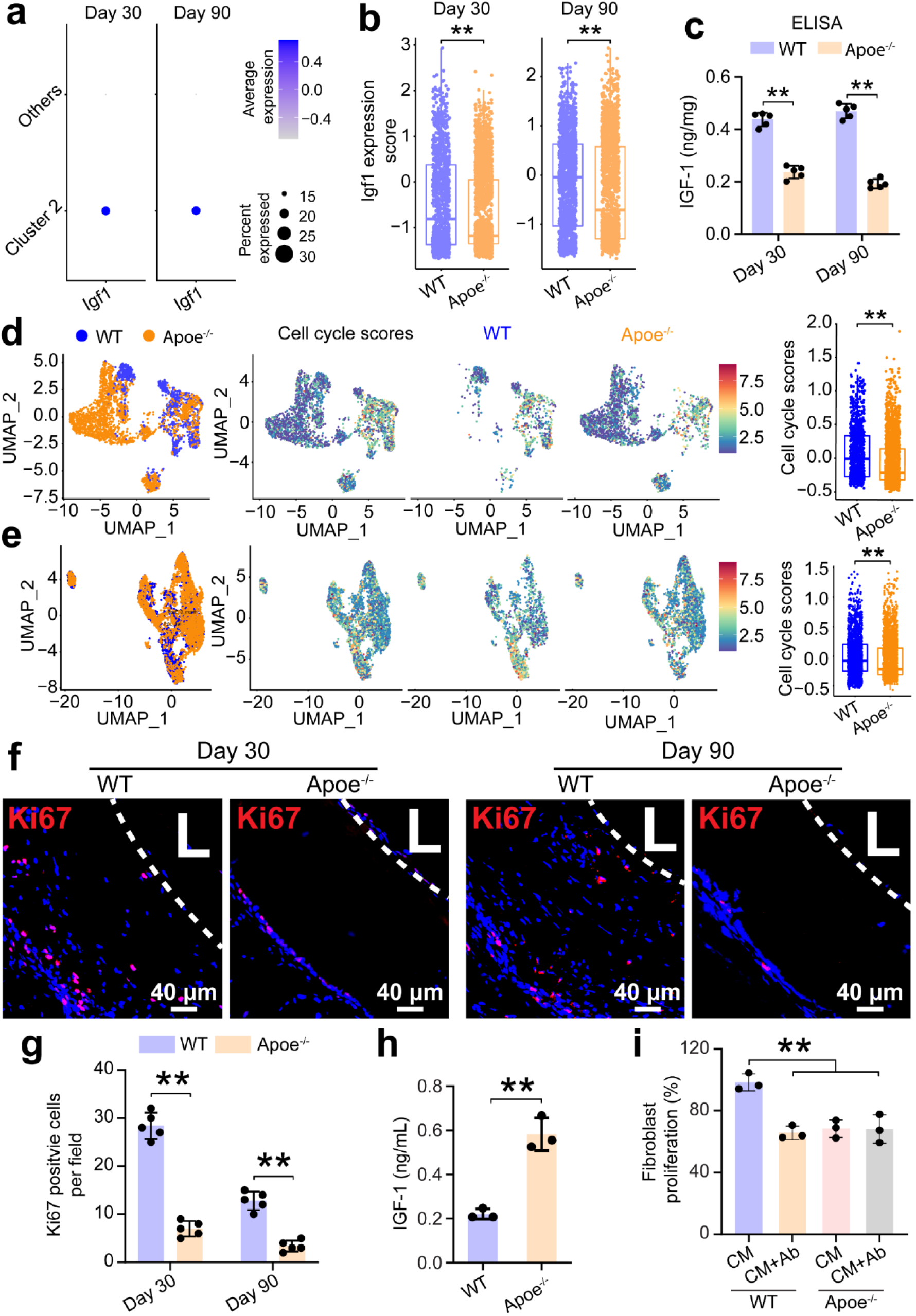
Fibroblast proliferation supported by profibrotic macrophages via IGF-1 secretion. Dot plots (a) and box plots (b) of Igf1 expression in macrophages of regenerated aortas post graft implantation in WT and Apoe^-/-^ rats on Day 30 and Day 90. (c) Quantification of IGF-1 by ELISA in regenerated aortas post graft implantation in WT and Apoe^-/-^ rats on Day 30 and Day 90. ** indicates p < 0.01, Tukey’s post-hoc test. For each timepoint and each group, five different samples from five different animals were quantified (n=5). UMAP of fibroblasts in regenerated aortas post graft implantation in WT and Apoe^-/-^ rats, heatmap of cell cycle scores in the UMAP of fibroblasts, and box plots of cell cycle scores in the fibroblasts on Day 30 (d) and Day 90 (e). ** indicates p < 0.01, unpaired t test. (f) Immunofluorescence staining of Ki67 in regenerated aortas post graft implantation in WT and Apoe^-/-^ rats on Day 30 and Day 90. L indicates lumens. (g) Quantification of Ki67 positive cells. ** indicates p < 0.01, Tukey’s post-hoc test. For each timepoint and each group, five different samples from five different animals were quantified (n=5). (h) Quantification of IGF-1 by ELISA in cell culture mediums after macrophages from WT or Apoe^-/-^ rats were cultured on PCL scaffolds for 48 hours. ** indicates p < 0.01, unpaired t test. For each timepoint and each group, three different samples were quantified (n=3). (i) Quantification of proliferation of fibroblasts treated with condition medium (CM) with or without IGF-1 blocking antibody (Ab, 1 µg/mL) for 24 hours using cell counting kit-8 (CCK-8). ** indicates p < 0.01, Tukey’s post-hoc test. For each timepoint and each group, three different samples from three different animals were quantified (n=3).

### Inhibition of APOE production attenuating fibrosis during vascular regeneration after graft implantation *in vivo*

To ameliorate fibrosis during vascular regeneration after graft implantation, adventitial delivery of adeno-associated virus (AAV) encoding shRNA targeting Apoe (AAV-shRNA(Apoe)) was used in this study to knockdown APOE *in vivo* (**Fig. 8a**). Two weeks after graft implantation *in vivo*, AAV-shRNA(Apoe) were injected into adventitia of regenerated aortas, which were then harvested for analysis three weeks later. Both *in vivo* imaging system (IVIS) and immunofluorescence imaging confirmed a successful transfection of cells in regenerated aortas by AAV (**Supplementary Fig. 9a**) and the transfection efficiency was around 30%-40% (**Supplementary Fig. 9b**). Ultrasound imaging showed that AAV virus treatment had no influence on graft patency (**Supplementary Fig. 9c**). However, movements of vascular walls could only be observed in grafts treated with AAV-shRNA(Apoe) but not in those treated with AAV-shRNA(NC) or phosphate buffered saline (PBS, the model group, **Fig. 8b**). After AAV-shRNA(Apoe) treatment, the regenerated aortas had a higher RI and PI than those treated with AAV-shRNA(NC) and PBS (**Fig. 8c**). More importantly, the compliance of the regenerated aortas was greatly improved compared with the AAV-shRNA(NC) and model groups. The AAV-shRNA(Apoe) group also had a thinner adventitia than the AAV-shRNA(NC) and model groups did, as shown in H&E staining in **Fig. 8d and g**. MTC staining revealed that there were less collagen deposition in the adventitial areas of the regenerated aortas in the AAV-shRNA(NC) group as compared with the AAV-shRNA(NC) and model groups as well (**Fig. 8d and h**). Meanwhile, APOE knockdown reduced the number of profibrotic macrophages (CD68^+^CTSD^+^) compared with the AAV-shRNA(NC) and model groups (**Fig. 8e and i**). Protein levels of APOE, CTSD and SPP1 in the AAV-shRNA(Apoe) group were also lower than those in the AAV-shRNA(NC) and model groups (**Fig. 8f and j**). The positive areas of IGF-1 after AAV-shRNA(Apoe) treatment also decreased as compared to the AAV-shRNA(NC) and model groups (**Supplementary Fig. 9d and e**). These results indicate that AAV targeting Apoe can successfully downregulate the expression of APOE during vascular regeneration after graft implantation *in vivo*, which further inhibited fibrosis by reducing profibrotic transition of macrophages and macrophage-derived IGF-1.

**Fig. 8.**
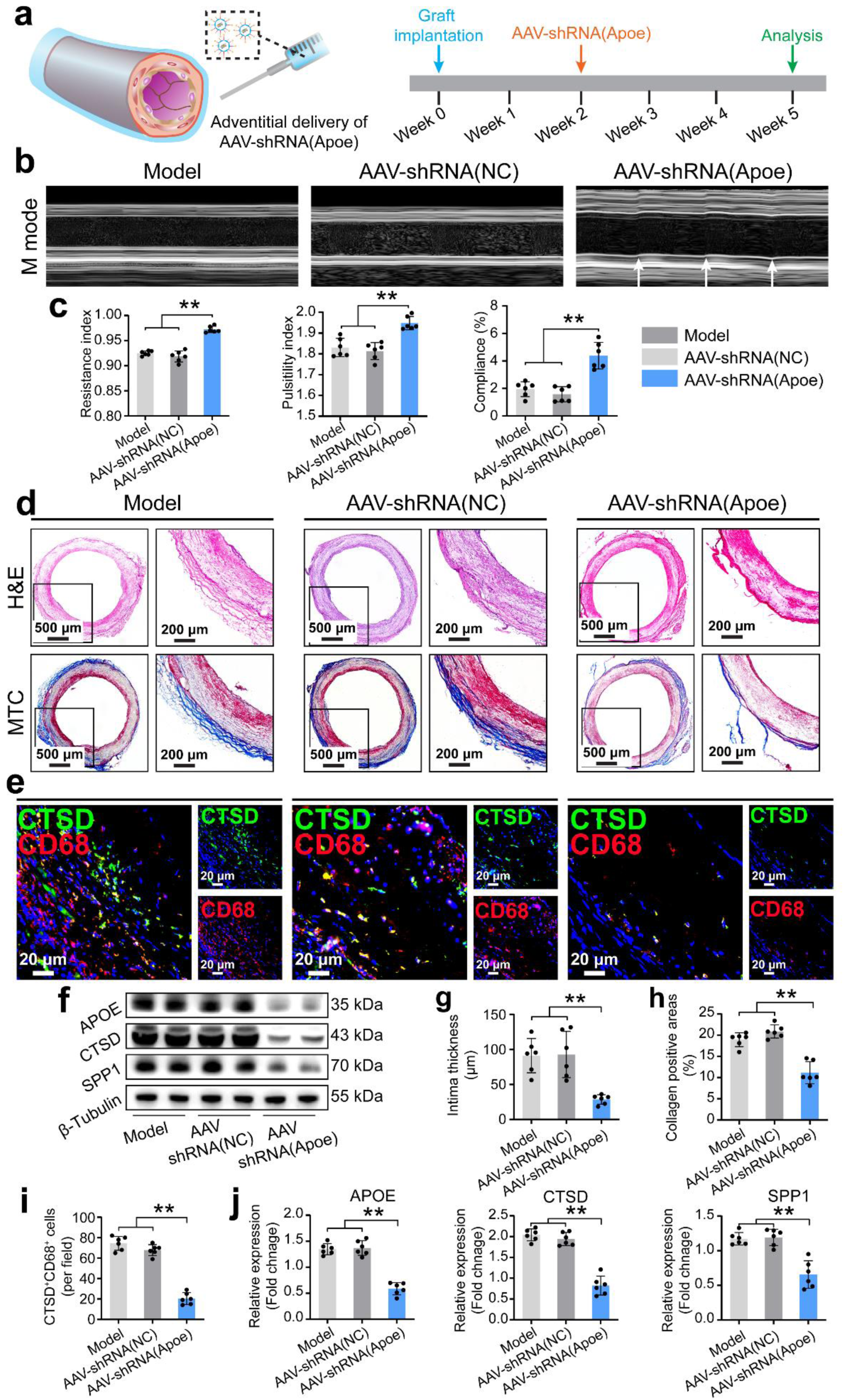
Downregulation of APOE by adventitial delivery of AAV ameliorating fibrosis during vascular regeneration after graft implantation. (a) Illustration of a strategy of adventitial delivery of AAV-shRNA(Apoe) to knockdown APOE. Two weeks after graft implantation *in vivo*, AAV-shRNA(Apoe) were injected into the adventitia of the regenerated aortas, which were then harvested for analysis three weeks later. (b) M mode images of ultrasound of regenerated aortas treated with PBS, AAV-shRNA(NC), and AAV-shRNA(Apoe). Arrow heads indicate movement of vascular walls. (c) Quantification of RI, PI, and compliance of regenerated aortas treated with PBS, AAV-shRNA(NC), and AAV-shRNA(Apoe). ** indicates p < 0.01, Tukey’s post-hoc test. For each group, six different images from six different animals were quantified (n=6). (d) H&E and MTC staining of regenerated aortas treated with PBS, AAV-shRNA(NC), and AAV-shRNA(Apoe). (e) Immunofluorescence staining of CTSD and CD68 in regenerated aortas treated with PBS, AAV-shRNA(NC), and AAV-shRNA(Apoe). (f) WB analysis of levels of APOE, CTSD and SPP1 in regenerated aortas treated with PBS, AAV-shRNA(NC), and AAV-shRNA(Apoe). Quantification of (g) intima thickness, (h) collagen positive areas, (i) CD68 and CTSD double positive cells, (j) levels of APOE, CTSD and SPP1 in regenerated aortas treated with PBS, AAV-shRNA(NC), and AAV-shRNA(Apoe). ** indicates p < 0.01, Tukey’s post-hoc test. For each group, six different samples from six different animals were quantified (n=6).

## Conclusion

In summary, we showed that APOE, which increased dramatically after vascular grafts implanted *in vivo*, induced profibrotic transition of macrophages via APOE/LRP1 interaction. Profibrotic macrophages increased fibroblast proliferation through IGF-1 to promote graft fibrosis. Inhibition of APOE by adventitial delivery of AAV targeting Apoe ameliorated graft fibrosis occurring during vascular regeneration. Our findings identify APOE as a novel therapeutic target to limit graft fibrosis during vascular regeneration.

## Methods

### PCL graft fabrication

Vascular grafts used in this study were made of polycaprolactone (PCL) via electrospinning as previously described with modifications^51^. Briefly, 14% PCL solutions were prepared by dissolving 14 g of PCL (80,000 in average molecular weight, Sigma-Aldrich) in 100 mL of 2,2,2-trifluoroethanol (Sigma-Aldrich). Then, the PCL solutions were pumped out from a nozzle at a rate of 2 mL per hour under a voltage of 12 kV. A 2-mm-thick stainless-steel mandrel rotating at a speed of 800 rpm was used to collect PCL fibers. The distance between the nozzle and mandrel was 15 cm. 600 μL of the 14% PCL solutions were used for each graft. Before the *in vivo* implantation, the grafts were cut into 10-mm-long segments, which were sterilized with 75% ethanol and dried in air.

### Animals

Animal experiments (Protocol No.: SRRSH202402504) were approved by the Chinese Institutional Animal Care and Use Committee at Sir Run Run Shaw Hospital, School of Medicine, Zhejiang University. Wide type (WT) SD rats used in this study were purchased from the Experimental Animal Center of Zhejiang Province. Apoe-knockout (KO, Apoe^-/-^) SD rats (SD-Apoe*^em1Smoc^*, NR-KO-190003) were purchased from the Shanghai Model Organisms Center. All animal experiments were performed on 8- to 10-week-old rats with a body weight around 400 g.

### Vascular graft implantation

Vascular regeneration was observed in a graft abdominal aorta interposition model in rats. Briefly, isoflurane was used to anesthetize rats, and the abdominal aortas were exposed after a midline incision on the abdomens. Then, the small branches of the abdominal aortas were ligated with 10-0 sutures, and the blood flow in the aortas was blocked with two microvascular clamps that were 1.5 cm apart. A 5-mm-long segment of the aortas above the renal arteries was removed, which was replaced by a 10-mm-long graft. End-to-end anastomosis at the two ends of the graft were performed with 8-0 sutures. At the end, the blood flow recovered after the release of the microvascular clamps, and the abdominal incision was closed with 4-0 sutures.

### Preparation of single cell suspension

For WT SD rats, after implantation of vascular grafts *in vivo* for 14, 30 and 90 days respectively, the regenerated abdominal aortas, as well as age matched native ones, were harvested for single cell suspension. For Apoe-KO SD rats, regenerated abdominal aortas were harvested for single cell suspension after vascular grafts were implanted *in vivo* for 30 and 90 days, respectively. Briefly, regenerated abdominal aortas or native aortas were rinsed with cold phosphate buffered saline (PBS) and cut into small pieces about 1×1 mm^2^ in size. Collagenases, hyaluronidases and dispases were used to dissociate the vascular tissues into single cells at 37 °C with shaking at a speed of 100 rpm for 45 minutes. The digested tissues were sequentially filtered through 70 μm and 40 μm cell strainers. After that, the filtered samples were treated with red blood cell lysis buffer (MeilunBio) at room temperature for 5 minutes. After centrifugation, cells resuspended in PBS, and cell viability and concentrations were determined using Countess II Automated Cell Counter (Thermo Fisher Scientific).

### Single cell transcriptome sequencing and data processing

When cell viability was higher than 85%, cell suspensions were adjusted to concentrations between 500 to 1500 cells per μL, and cell suspensions were then loaded into Chromium controller (10x Genomics) for single cell partitions, molecular barcode labeling and cDNA synthesis as instruction manuals. Finally, the constructed cDNA was pooled, amplified, and sequenced. The above sequencing work were finished by LC-Bio Technology (Hangzhou, China). The generated single cell transcriptome data were processed using online tools (https://www.omicstudio.cn) provided by LC-Bio Technology. CellPhoneDB module of the online tools (v5.0.0) was used to analyze communications among different types of cells.

### Immunofluorescence staining

After sample harvest, 4% paraformaldehyde solution was used to fix the explanted samples at 4 °C for 1 hour, and then 30% sucrose solution was used for sample dehydration at 4 °C overnight. Then, samples were sectioned at a thickness of 8 μm after they were embedded in O.C.T. compound (Tissue-Tek) and frozen at -80 °C. PBS was used to wash tissue sections and 5% bovine serum albumin (BSA, Sigma-Aldrich) solution was used for background blockage at room temperature for 30 minutes. Primary antibodies were diluted with 5% BSA solution first and then used to incubate samples at 4 °C overnight. The following primary antibodies were used in this study: APOE (Invitrogen, PA5-78803, 1:200 dilution), COL I (abcam, ab270993, 1:200 dilution), COL III (abcam, ab6310, 1:200 dilution), LUM (abcam, ab252925, 1:200 dilution), CTSD (CST, 74089S, 1:200 dilution), CD68 (BioRad, MCA341GA, 1:100 dilution), LRP1 (Invitrogen, PA5-101013, 1:200 dilution), Ki67 (Servicebio, GB111141, 1:200 dilution), and IGF1 (Invitrogen, MA5-18035, 1:200 dilution). After that, PBS was used to wash tissue sections again and secondary antibodies (Invitrogen) were used to incubate the samples at room temperature for 1 hour. Tissue sections incubated with secondary antibodies but not with primary antibodies were used as negative controls to rule out false positive staining. Nuclei were counterstained with 1 μg/mL 4′,6-Diamidino-2-phenylindole dihydrochloride (DAPI, Sigma-Aldrich). Finally, both stained samples and negative controls were observed with an inverted fluorescence microscope (Olympus) with the same imaging settings.

### Western blot (WB)

RIPA buffer (Solarbio) with 1% protease inhibitor PMSF (Solarbio) were used to extract proteins from samples. BCA Protein Assay Kits (Thermo Fisher Scientific) were used to quantify protein concentrations of the extracted proteins, and then 5×loading buffer (Fude Biological Technology) was used to denature the protein samples by heating to 95 °C for 5 minutes. 4-20% sodium dodecyl sulfate-polyacrylamide electrophoresis gel (ACE Biotechnology) was used to separate proteins into different bands, which were transferred to polyvinylidene difluoride (PVDF) membranes (Millipore) for antibody detection. 5% non-fat milk was used for background blockage at room temperature for 1 hour, and then diluted primary antibodies were used to incubate samples at 4 °C overnight. The following primary antibodies were used in this study: APOE (Abcam, ab183597, 1:1000 dilution), COL I (abcam, ab270993, 1:1000 dilution), FN (abcam, ab268020, 1:1000 dilution), CTSD (CST, 74089S, 1:1000 dilution), SPP1 (NeoBiotechnologies, 6696-RBM3-P0, 1:1000 dilution), and LRP1 (Invitrogen, PA5-101013, 1:200 dilution). After that, samples were further incubated in diluted horseradish peroxidase (HRP)-conjugated secondary antibodies (Fude Biological Technology) at room temperature for 1 hour. The expression of β-Tubulin was detected with β-Tubulin-HRP (Proteintech, 66240-1-Ig, 1:5000 dilution) as an internal control. Enhanced chemiluminescence kit (ABclonal) was used to treat samples before they were imaged with Amersham Imager 600 (GE, UK).

### Enzyme linked immunosorbent assay (ELISA)

Samples were collected, rinsed with cold PBS, weighed, and minced. A tissue homogenizer was used to homogenize tissues at 4 °C for 10 minutes. After centrifugation at a speed of 5000×g at 4 °C for 10 minutes, supernatant was collected for ELISA. For cell culture medium, after centrifugation at a speed of 5000×g at 4 °C for 20 minutes, supernatant was collected for ELISA. An APOE ELISA kit (Ruixin Biotech, RX302092R) or IGF-1 ELISA kit (Ruixin Biotech, RX302143R) was used as manuals.

### Graft performance evaluation *in vivo* by ultrasound imaging

A small animal ultrasound imaging system (VisualSonics, Vevo 3100, FUJIFILM) was used to evaluate graft performance in WT and Apoe-KO SD rats after implantation *in vivo* for 30 or 90 days, respectively. Briefly, isoflurane was used to anesthetize rats, and abdominal aortas were exposed via an incision made on abdomens. The ultrasound probes touched the abdominal aortas directly to acquire B mode, color mode, PW mode, and M mode images. The resistance index (RI) and pulsatility index (PI) were calculated from the images of M mode using software provided by FUJIFILM. The compliance of the blood vessels was calculated from the images of B mode in long axis.

### Chemical staining

Samples were harvested, fixed, dehydrated, and sectioned as aforementioned. Tissue sections were first washed with PBS and then stained with hematoxylin and eosin (H&E, Servicebio) and Masson’s trichrome (MTC, Solarbio) using commercial kits as manuals. An inverted microscope (Olympus) was used to observe the stained samples.

### Subcutaneous implantation model

To fabricate PCL sheets for subcutaneous implantation, a 10-mm-thick stainless-steel mandrel rotating at a speed of 800 rpm was used to collect PCL fibers. The distance between the nozzle and mandrel was 30 cm. 800 μL of the 14% PCL solutions were used for each sheet. Before the *in vivo* implantation, the PCL sheets were cut into small pieces with a size of 1×1 cm^2^, which were sterilized with 75% ethanol and dried in air. A pocket was made under the skin of the abdomens of rats, and a piece of PCL sheet was inserted into the pocket. After PCL sheets were subcutaneously implanted *in vivo* for 30 or 90 days in WT and Apoe-KO SD rats respectively, samples were harvested, fixed, dehydrated, and sectioned for histological evaluation as aforementioned.

### Fibroblast isolation and culture

Thoracic aortas and the surrounding tissues were collected from WT SD rats, which were further washed with PBS. Adventitial tissues were separated from the thoracic aortas and minced into small pieces, which were seeded onto bottoms of flasks. 2 hours later, Dulbecco’s Modified Eagle Medium (DMEM, Gibico) with 10% fetal bovine serum (FBS, Biological Industries) was added into the flasks and the minced tissues were cultured in a humidified environment with 5% CO_2_ at 37 °C. Outgrowth cells surrounding tissues were digested with trypsin, resuspended in DMEM with 10% FBS, and plated in a new flask. Fibroblasts with a passage number 3 to 8 were used in this study. Exogenous APOE (MCE, HY-P701096), conditioned medium by macrophages, or IGF-1 blocking antibody (Invitrogen, MA5-18035) was added into culture medium as indicated.

### Peritoneal macrophage isolation and culture

Peritoneal macrophages were extracted from abdominal cavities of WT or APOE KO SD rats after starch stimulation. Briefly, the abdominal cavities of rats were injected with 6% sterile starch solution (Solarbio) 3 days before macrophage extraction. After that, the abdominal cavities of the rats were first flushed with 40 mL of PBS, and then the lavage fluid were centrifuged at 1000 rpm at room temperature for 10 minutes to pellet the macrophages. The isolated macrophages were resuspended in DMEM with 10% FBS and seeded on 1×1 cm^2^ PCL sheets, which were cultured in a humidified environment with 5% CO_2_ at 37 °C for 48 hours. As a control, the isolated macrophages were seeded onto tissue culture plates and cultured under the same conditions.

### Immunocytochemistry

Cells were washed with PBS for thrice, fixed in 4% paraformaldehyde at room temperature for 10 minutes, and then permeabilized with 0.1% Triton X-100 (Fude Biological Technology) at room temperature for another 10-minute period. Then, the cells were incubated with primary antibodies diluted with 5% BSA at room temperature for 1 hour. The following primary antibodies were used in this study: APOE (Invitrogen, 701241, 1:200 dilution), CTSD (CST, 74089S, 1:200 dilution), SPP1 (NeoBiotechnologies, 6696-RBM3-P0, 1:200 dilution), CD68 (BioRad, MCA1957GA, 1:100 dilution), and LRP1 (Invitrogen, PA5-101013, 1:200 dilution). After that, PBS was used to wash cells again, and the secondary antibodies were used to incubate the cells at room temperature for 1 hour. 1 μg/mL DAPI was used to counterstain nuclei, and the stained samples were imaged with a confocal microscope (LSM 800, Zeiss).

### Immunoprecipitation (IP) and mass spectrometry (MS)

To identify the receptors of APOE in macrophages, immunoprecipitation (IP) was performed followed by mass spectrometry (MS). Briefly, peritoneal macrophages were treated with PBS or 0.5 μg/mL APOE (MCE, HY-P701096) for 24 hours. IP lysis buffer (Pierce, Thermo Fisher Scientific) was used to lyse cells on ice for 30 minutes. Supernatants were collected for subsequent analysis after centrifugation at a speed of 1000 rpm at 4 °C. Protein A/G magnetic beads (MCE) were incubated with the APOE antibody (Abcam, ab183597, 1:40 dilution) at 4 °C overnight. Then, the supernatants of cell lysates were incubated with the coated magnetic beads at 4 °C overnight. After that, magnetic stands were used to isolate the magnetic beads, which were mixed with 5×loading buffer and heated to 95 °C for denaturation before electrophoresis. Finally, proteins were extracted from gel strips, and MS was performed by LC-Bio Technology to identify proteins that could possibly interact with APOE.

### Co-immunoprecipitation (Co-IP)

To confirm proteins that could directly interact with APOE, co-immunoprecipitation (Co-IP) was performed. Briefly, peritoneal macrophages were treated with PBS or 0.5 μg/mL APOE for 24 hours. After that, cells were lysed, and supernatants were collected as aforementioned. 10% of the supernatants were used as Input. Protein A/G magnetic beads (MCE) were coated with the APOE antibody as aforementioned, which were used to pull down APOE and proteins bound to APOE. The Input and proteins pulled down by the magnetic beads were detected using WB as aforementioned.

### Transfection of macrophages with adenovirus (ADV)

To knock down Lrp1 expressions in macrophages, adenovirus (ADV, OBiO) was used in this study. Vectors of pADV-U6-shRNA(Lrp1)-CMV-EGFP and pADV-U6-shRNA(NC2)-CMV-EGFP were used to generate ADV-shRNA (Lrp1) and ADV-shRNA (NC), respectively. Transfection efficiency was determined according to percentage of cells expressing EGFP. When the multiplicity of infection (MOI) was 240, the transfection efficiency was higher than 80%. Therefore, 6.75 μL of 7.11×10^10^ PFU/mL ADV-shRNA (Lrp1) or 2 μL of 2.37×10^11^ PFU/mL ADV-shRNA (NC) was used to transfect 2 million cells to ensure a MOI of 240. 24 hours later, the transfected cells were dissociated with trypsin, resuspended in DMEM with 10% FBS and seeded on 1×1 cm^2^ PCL scaffolds, which were cultured in a humidified environment with 5% CO_2_ at 37 °C for another 48-hour period.

### Flow cytometry

Trypsin was used to dissociate cells from PCL scaffolds, which were then washed with PBS, fixed in 4% paraformaldehyde at room temperature for 10 minutes, and then permeabilized with 0.1% Triton X-100 at room temperature for another 10-minute period. Then, the cells were incubated with CTSD (CST, 74089S, 1:200 dilution) at room temperature for 1 hour. After that, PBS was used to wash cells again, and the secondary antibodies were used to incubate the cells at room temperature for 1 hour. Samples were analyzed using flow cytometer (BD, FACSCalibur).

### Adventitial delivery of adeno-associated virus (AAV)

To inhibit excessive production of APOE during vascular regeneration, adventitial delivery of adeno-associated virus (AAV, OBiO) was used in this study as previously reported^52,53^. Vectors of pAAV-U6-shRNA(Apoe)-CMV-mScarlet-WPRE and pAAV-U6-shRNA(NC)-CMV-mScarlet-WPRE were used to generate AAV-shRNA (Apoe) and AAV-shRNA (NC), respectively. The serotype of AAV was AAV2/5. 14 days after graft implantation *in vivo*, In the AAV treated groups, 100 μL of 4.74×10^12^ v.g./mL AAV-shRNA (Apoe) or 100 μL of 3.89×10^12^ v.g./mL AAV-shRNA (NC) was injected into the adventitial areas of the regenerated abdominal aortas. In the model group, 100 μL of PBS was injected. 21 days later, the performance of vascular grafts *in vivo*, the vascular graft histology and the expressions of target proteins in vascular grafts were evaluated.

## Data analysis

All data were presented as mean ± standard deviation. Each test had at least three replicates and repeated thrice independently. The unpaired t-test was used for two-group comparison, and the one-way ANOVA followed by Tukey’s post-hoc test was used to analyze the data. P < 0.05 was statistically significant in this study.

## Supporting Information

Supplementary Figures.

## Acknowledgments

This work was sponsored by the Zhejiang Province Natural Science Foundation of China (LQ22H180001), the Science and Technology of Medicine and Health program of Zhejiang Province (2023RC028, 2024KY102), the Natural Science Foundation of China (82300565), the Science and Technology project of Zhejiang Province (2024C03025), and the Traditional Chinese Medicine Modernization projects of Zhejiang Province (2022ZX012).

## Supplementary Figures

**Supplementary Fig. 1.**
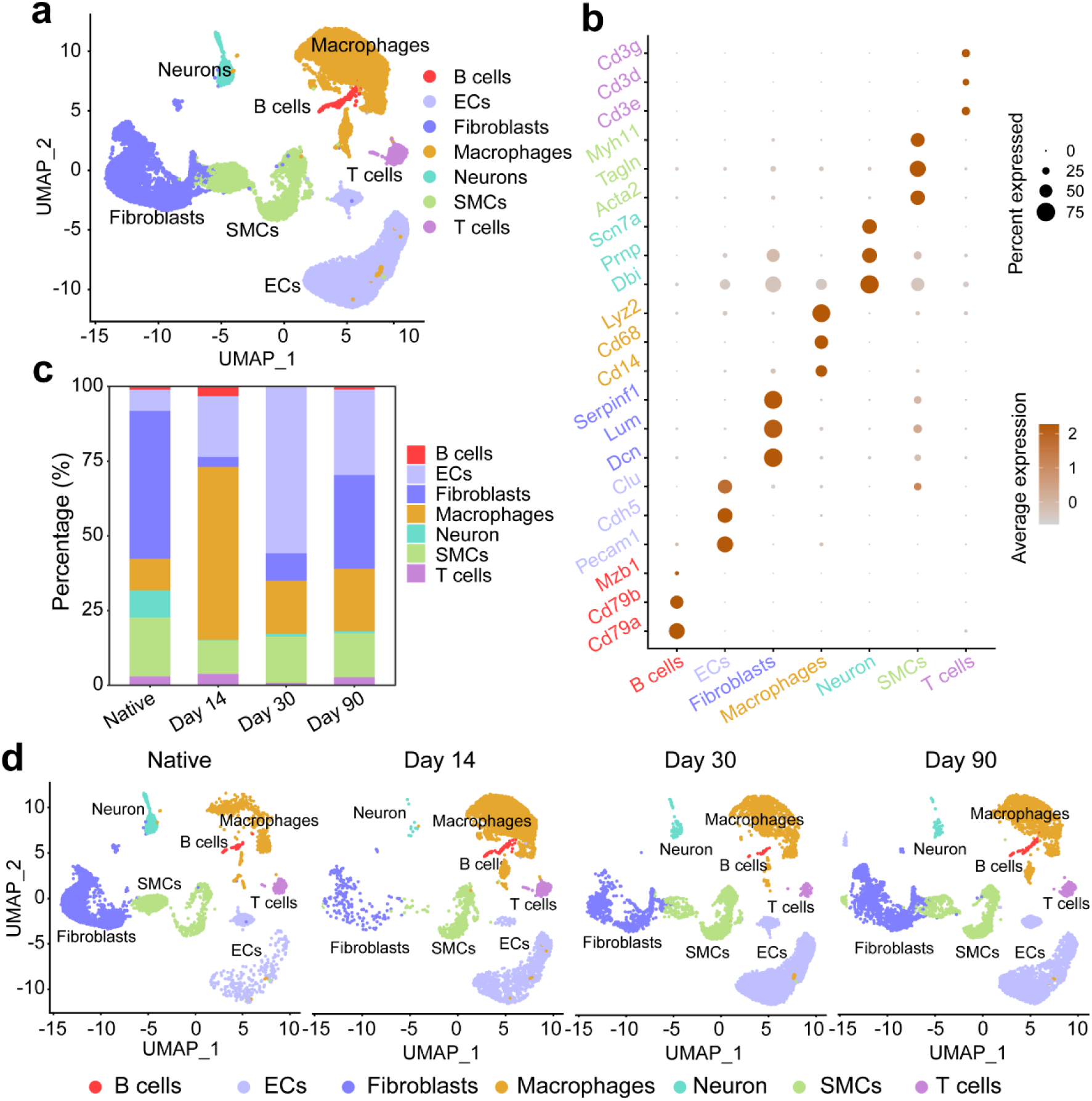
Single cell RNA sequencing (scRNA-seq) of native aortas and regenerated aortas 14, 30, and 90 days after vascular graft implantation in rats. (a) UMAP of different cell clusters in native aortas and regenerated aortas. (b) Dot plots of marker genes for different cell clusters. (c) Percentage of different cell clusters in native aortas and regenerated aortas across different timepoints post graft implantation. (d) UMAP of different cell clusters in native aortas and regenerated aortas across different timepoints post graft implantation.

**Supplementary Fig. 2.**
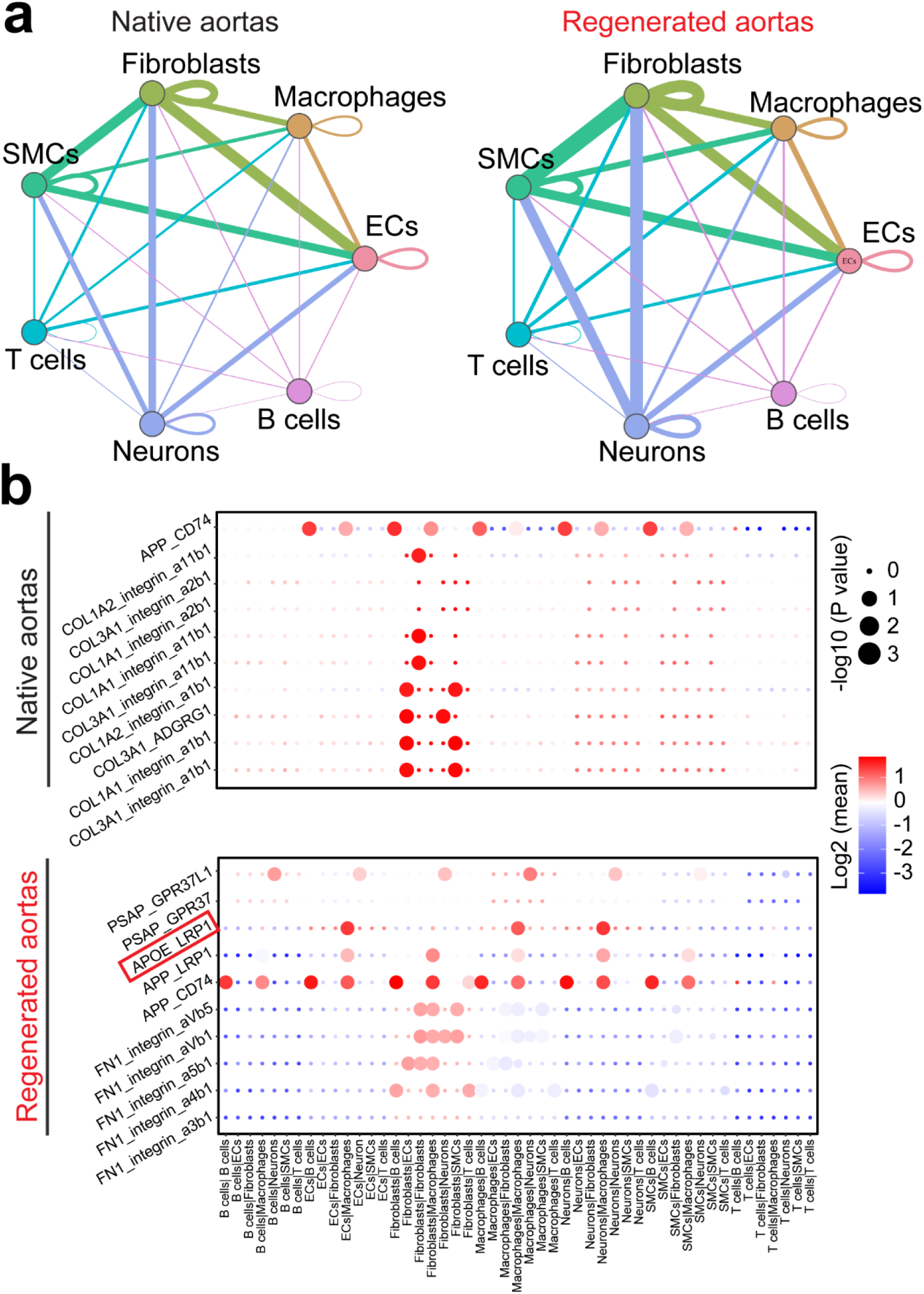
Crosstalk analysis between different cell types in native aortas and regenerated aortas by CellPhoneDB module. (a) Crosstalk networks in native aortas and regenerated aortas. The thickness of line between different cell clusters indicates number of ligand-receptor pairs. (b) Top 10 ligand-receptor pairs in native arteries and regenerated aortas.

**Supplementary Fig. 3.**
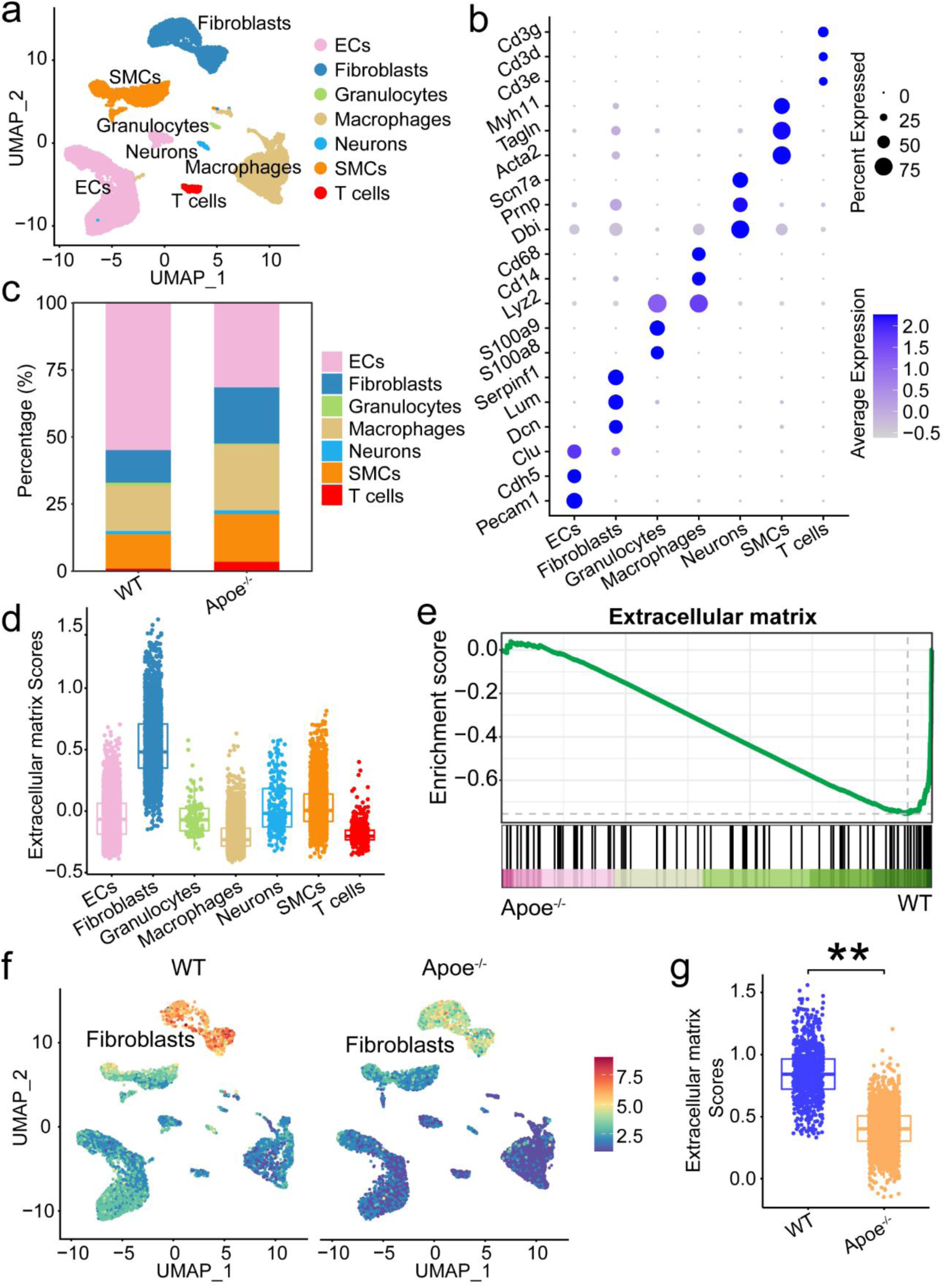
scRNA-seq of regenerated aortas from WT and Apoe^-/-^ rats at Day 30. (a) UMAP of different cell clusters in regenerated aortas from WT and Apoe^-/-^ rats. (b) Dot plots of marker genes for different cell clusters in regenerated aortas from WT and Apoe^-/-^ rats. (c) Percentage of different cell clusters in regenerated aortas from WT and Apoe^-/-^ rats. (d) Box plots of extracellular matrix scores across different cell clusters in regenerated aortas from WT and Apoe^-/-^ rats. (e) Gene set enrichment analysis (GSEA) of expressions of extracellular matrix-related genes by fibroblasts in regenerated aortas from WT and Apoe^-/-^ rats. (f) Heatmap of extracellular matrix scores across different cell clusters in regenerated aortas from WT and Apoe^-/-^ rats. (g) Box plots of extracellular matrix scores in fibroblasts in regenerated aortas from WT and Apoe^-/-^ rats. ** indicates p < 0.01, unpaired t test.

**Supplementary Fig. 4.**
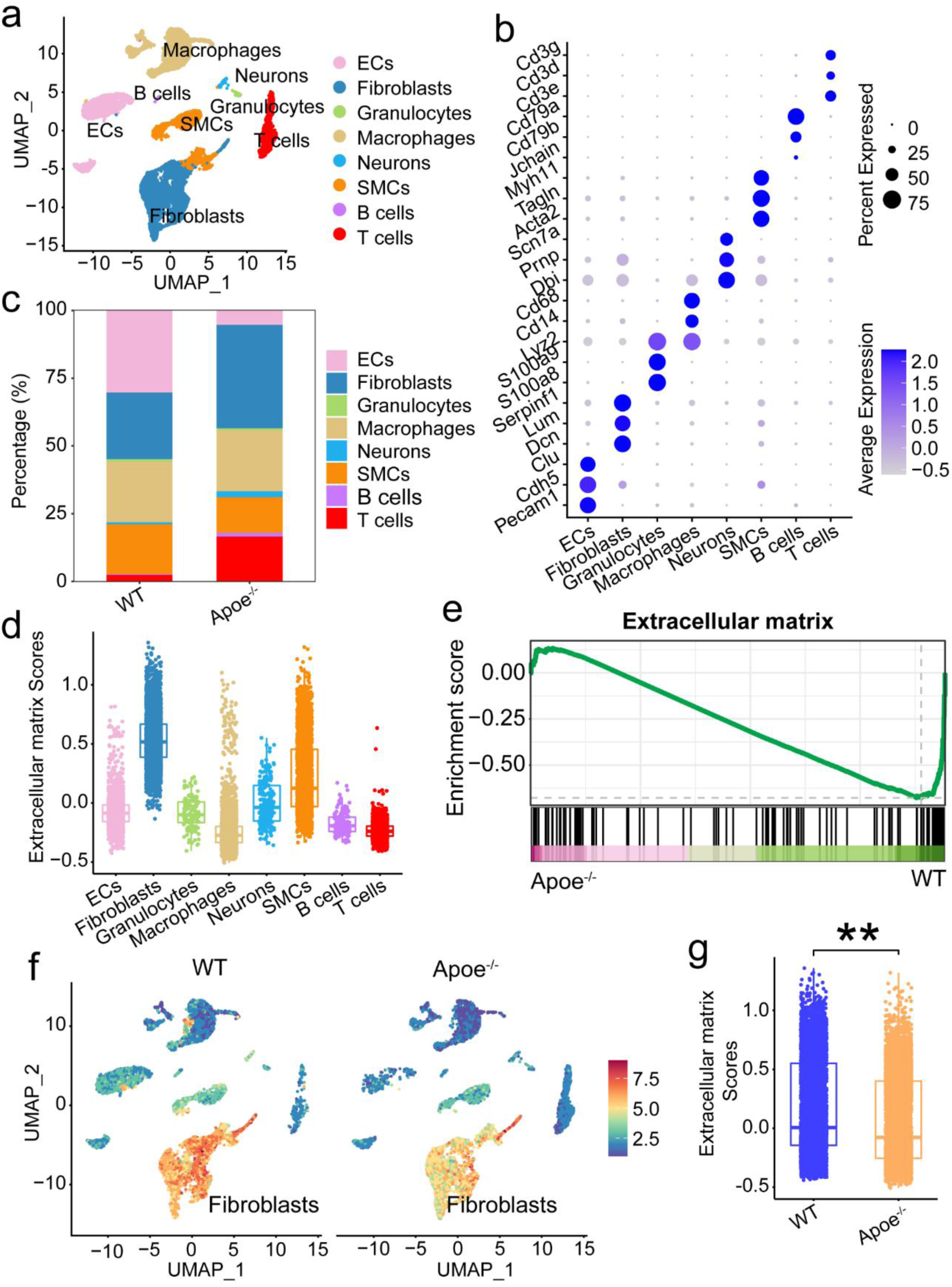
scRNA-seq of regenerated aortas from WT and Apoe^-/-^ rats at Day 90. (a) UMAP of different cell clusters in regenerated aortas from WT and Apoe^-/-^ rats. (b) Dot plots of marker genes for different cell clusters in regenerated aortas from WT and Apoe^-/-^ rats. (c) Percentage of different cell clusters in regenerated aortas from WT and Apoe^-/-^ rats. (d) Box plots of extracellular matrix scores across different cell clusters in regenerated aortas from WT and Apoe^-/-^ rats. (e) Gene set enrichment analysis (GSEA) of expressions of extracellular matrix-related genes by fibroblasts in regenerated aortas from WT and Apoe^-/-^ rats. (f) Heatmap of extracellular matrix scores across different cell clusters in regenerated aortas from WT and Apoe^-/-^ rats. (g) Box plots of extracellular matrix scores in fibroblasts in regenerated aortas from WT and Apoe^-/-^ rats. ** indicates p < 0.01, unpaired t test.

**Supplementary Fig. 5.**
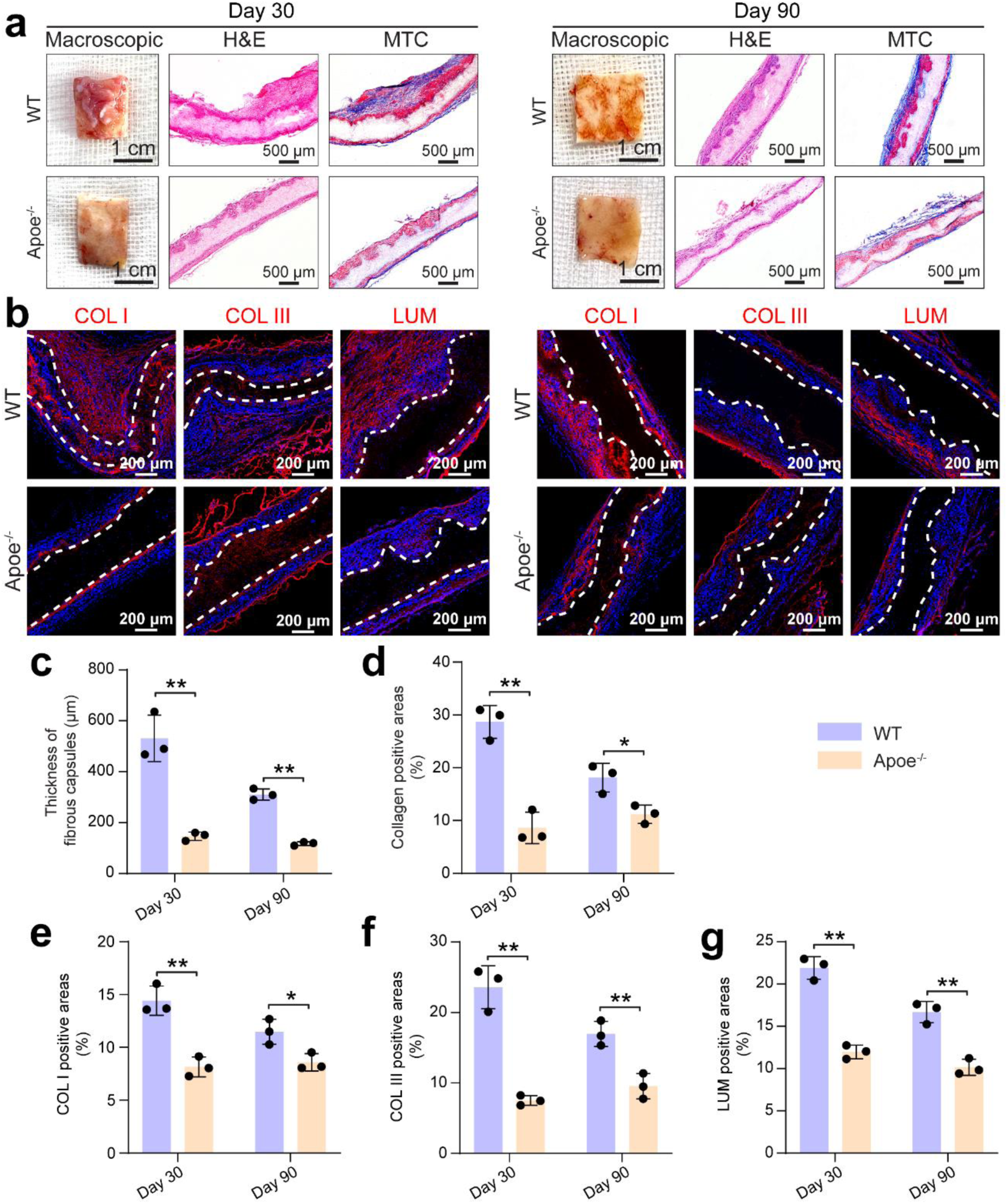
APOE KO limiting fibrous capsule formation of subcutaneously implanted PCL scaffolds. (a) Macroscopic views, H&E, and MTC staining of PCL scaffolds subcutaneously implanted in WT and Apoe^-/-^ rats at Day 30 and Day 90. (b) Immunofluorescence staining of COL I, COL III and LUM in PCL scaffolds subcutaneously implanted in WT and Apoe^-/-^ rats at Day 30 and Day 90. Areas between two dash lines indicate PCL scaffolds. Quantification of thickness of fibrous capsules (c), collagen positive areas (d), COL I positive areas (e), COL III positive areas (f), and LUM positive areas (g) in PCL scaffolds subcutaneously implanted in WT and Apoe^-/-^ rats at Day 30 and Day 90. ** indicates p < 0.01, Tukey’s post-hoc test. For each timepoint and each group, three different samples from three different animals were used (n=3).

**Supplementary Fig. 6.**
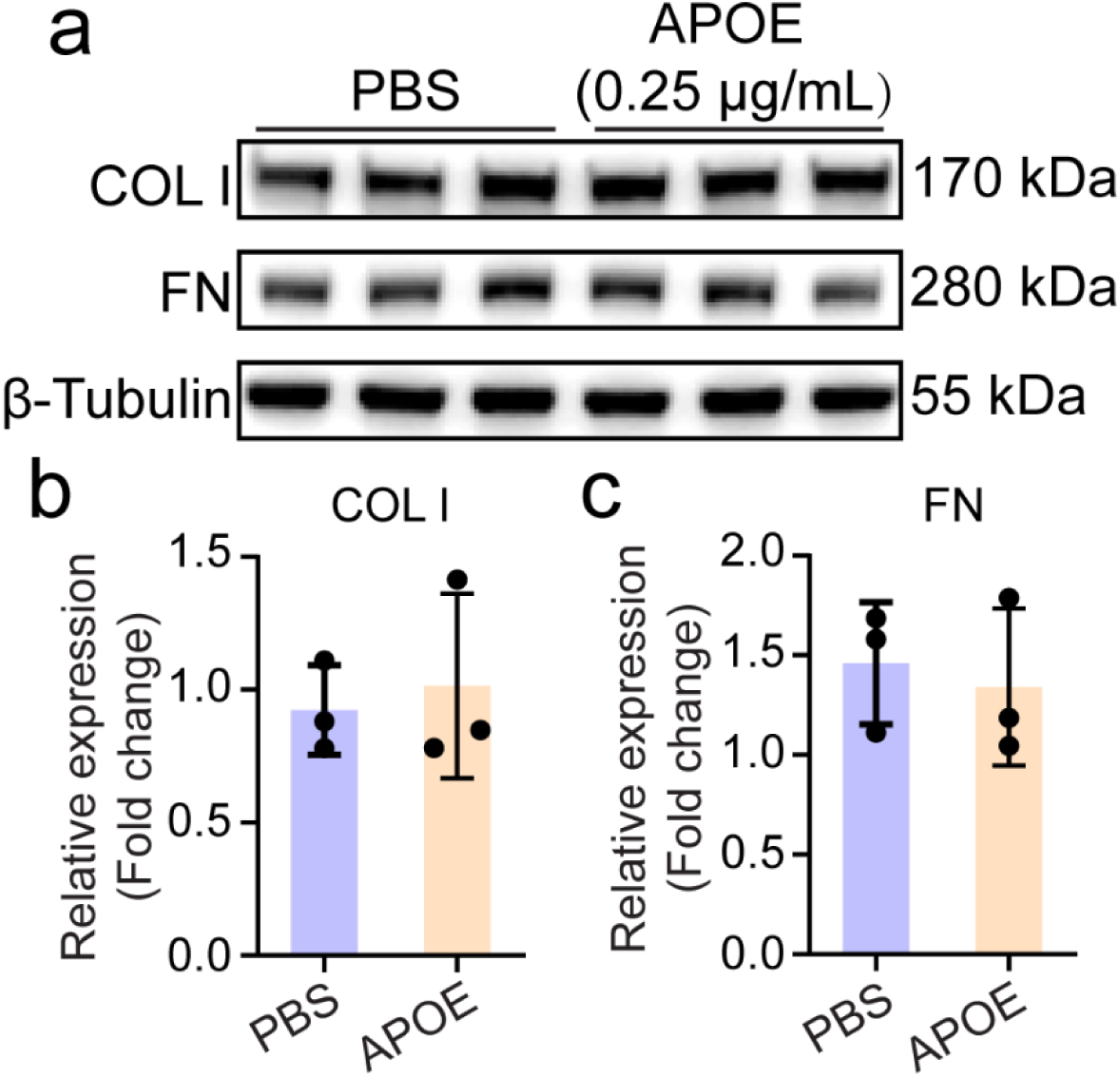
APOE treatment had no effects on ECM production by fibroblasts. (a) WB analysis of COL I and FN levels after treatment of fibroblasts with PBS or exogenous APOE (0.25 μg/mL) for 24 hours. (b) Quantification of COL I and FN levels. No significant difference is detected using unpaired t test. For each timepoint, three different samples were quantified (n=3).

**Supplementary Fig. 7.**
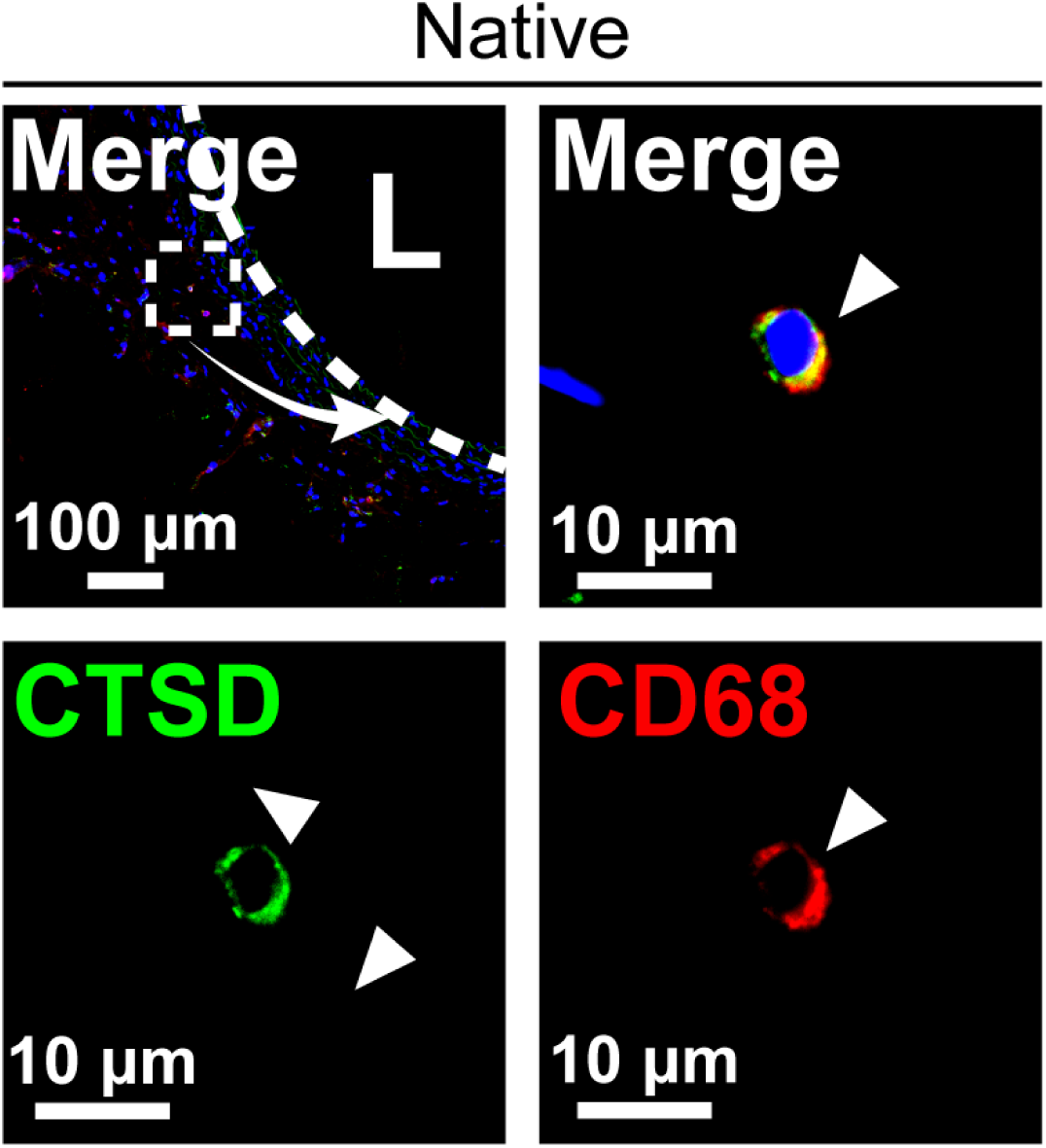
Immunofluorescence staining of CD68 and CTSD in native aortas. L indicates lumens. Arrow heads indicate positively stained cells.

**Supplementary Fig. 8.**
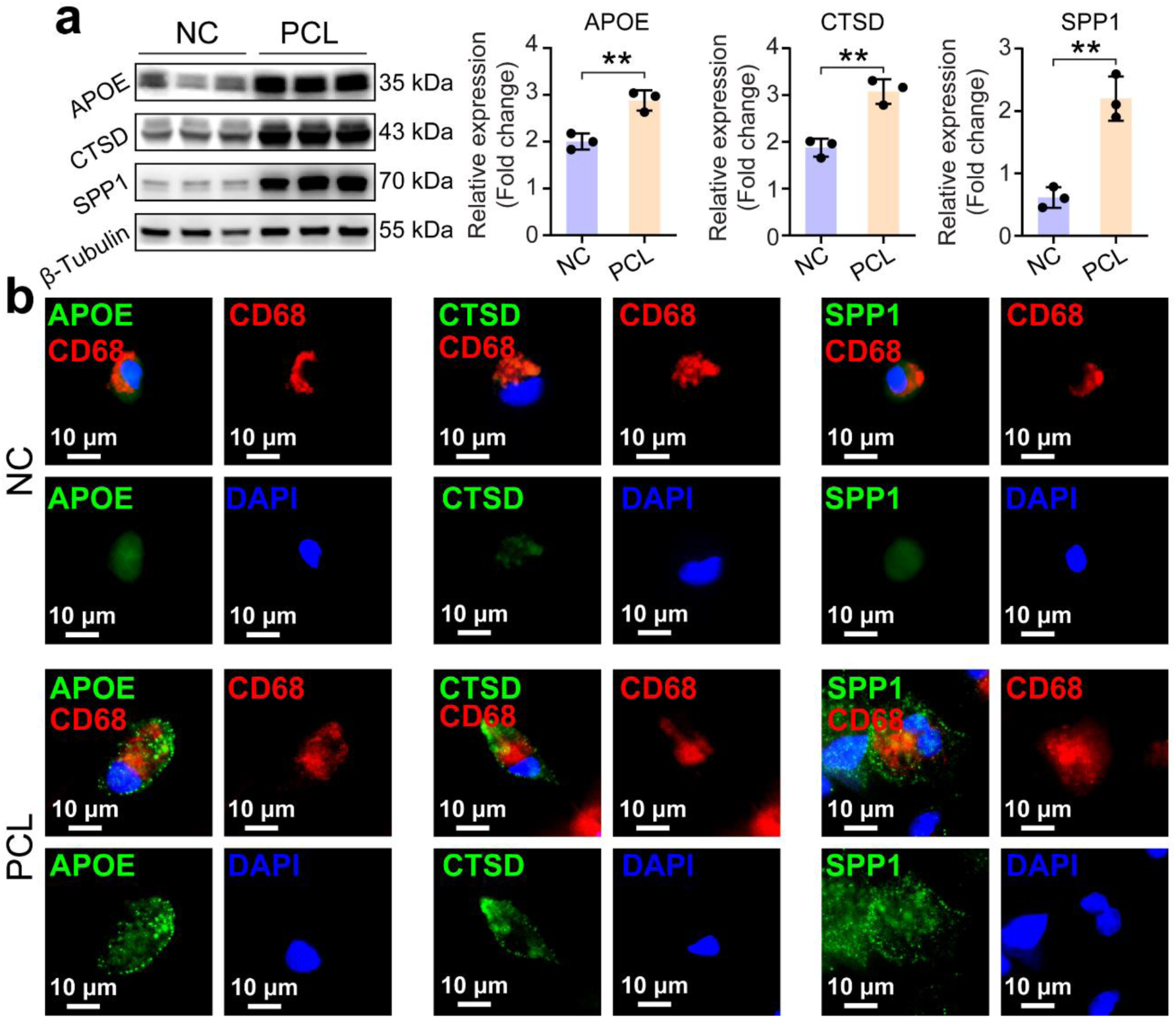
(a) WB analysis of levels of APOE, CTSD and SPP1 in macrophages cultured on tissue culture plates (negative control, NC) and PCL scaffolds (PCL) for 48 hours and quantification of their levels. ** indicates p < 0.01, unpaired t test. For each timepoint and each group, three different samples were quantified (n=3). (b) Immunofluorescence staining of APOE and CD68, CTSD and CD68, SPP1 and CD68 in macrophages cultured on tissue culture plates and PCL scaffolds.

**Supplementary Fig. 9.**
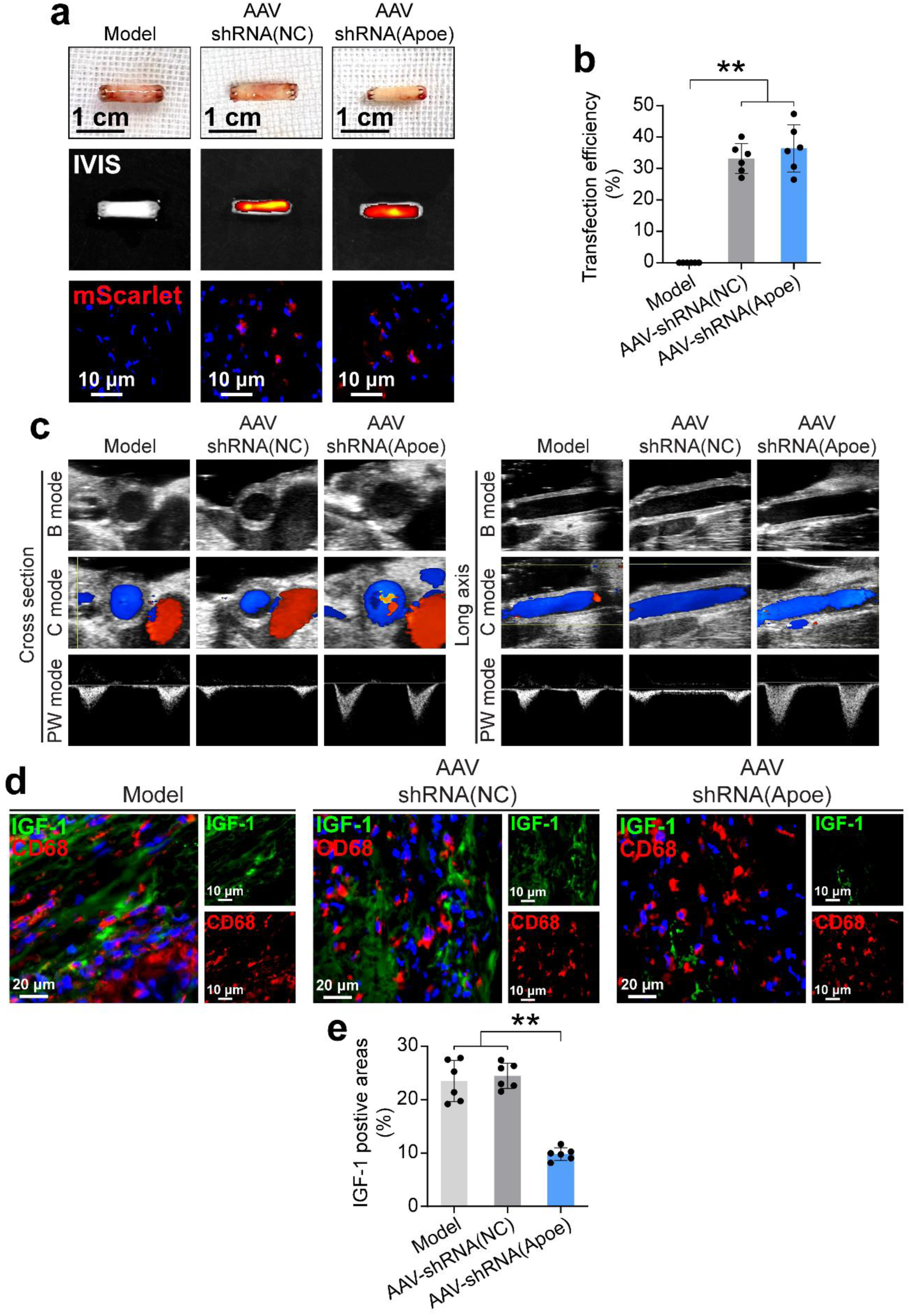
(a) Macroscopic views, *in vivo* imaging system (IVIS), and immunofluorescence imaging of regenerated aortas treated with PBS, AAV-shRNA(NC), and AAV-shRNA(Apoe). (b) Quantification of transfection efficiency of regenerated aortas by AAV according to mScarlet expression. ** indicates p < 0.01, Tukey’s post-hoc test. For each group, six different images from six different animals were quantified (n=6). (c) Ultrasound imaging of regenerated aortas treated with PBS, AAV-shRNA(NC), and AAV-shRNA(Apoe). (d) Immunofluorescence staining of IGF-1 and CD68 in regenerated aortas treated with PBS, AAV-shRNA(NC), and AAV-shRNA(Apoe). (e) Quantification of IGF-1 positive areas in regenerated aortas treated with PBS, AAV-shRNA(NC), and AAV-shRNA(Apoe). ** indicates p < 0.01, Tukey’s post-hoc test. For each group, six different samples from six different animals were quantified (n=6).

